# Visuomotor mismatch EEG responses over occipital cortex of freely moving human subjects

**DOI:** 10.1101/2025.08.14.670295

**Authors:** Magdalena Solyga, Marek Zelechowski, Georg B. Keller

## Abstract

Likely the strongest predictor of visual feedback is self-motion. In mice, the coupling between movement and visual feedback is learned with first visual experience of the world (Attinger et al., 2017), and brief perturbations of the coupling result in strong visuomotor mismatch responses in visual cortex that possibly reflect prediction errors (Keller et al., 2012; Zmarz and Keller, 2016). In humans, predictive coding has primarily been studied using oddball paradigms which rely on violations of stimulus probability based on recent sensory history. It was still unclear, however, whether humans exhibit visuomotor mismatch responses similar to those observed in mice. This question was important for two reasons. First, visuomotor mismatch responses in humans constitute a basis to start translating the mechanistic understanding of the circuit that computes these responses from mouse to human cortex. Second, a paradigm that can trigger strong prediction error responses and consequently requires shorter recording times would simplify experiments in a clinical setting. Here, by combining a wireless EEG recording system with a virtual reality headset, we found robust visuomotor mismatch responses in human cortex that were characterized by a reversed polarity relative to visual evoked responses and a greater signal power than both visual responses and oddball mismatch responses.

## INTRODUCTION

Predictive processing is a framework for understanding brain function. It proposes that the brain constructs internal models of the world based on regularities in past sensory input and sensorimotor loops to predict upcoming sensory input. The brain does this by minimizing the mismatches between predicted and actual sensory input. These mismatches, known as prediction errors, play a dual role in both updating internal representations and internal models (Friston, 2005; Keller and Sterzer, 2024; Rao and Ballard, 1999).

Predictions about sensory input can be made based on a variety of sources of other information the brain has available. Certain stimuli can be predicted from the preceding sensory input – experimentally this can be used, e.g., in oddball paradigms to trigger stimulus history based prediction errors (Garrido et al., 2009). Predictions can be crossmodal. Certain sounds are associated with specific types of visual input. Experimentally, visual stimuli can be coupled to sounds to then trigger audiovisual prediction errors in mouse visual cortex (Garner and Keller, 2022). Predictions can also be based on spatial memory. Not seeing a stimulus in a specific spatial location can result in visuospatial prediction errors in mouse cortex (Fiser et al., 2016). However, the strongest predictor - by frequency of experience, consistency in coupling, and precision in timing - is self-generated movement. Every movement is directly coupled to sensory feedback throughout life. Almost all visual and proprioceptive sensory input is the consequence of self-motion.

An essential ingredient to the computation of visuomotor prediction error responses are motor-related predictions of visual feedback (Keller and Mrsic-Flogel, 2018). In the mouse, evidence for such predictions has come from the discovery of a strong motor-related drive of activity in visual cortex (Keller et al., 2012; Niell and Stryker, 2010; Saleem et al., 2013). These motor-related signals are likely in part driven by motor-related predictions arriving from motor areas of cortex (Leinweber et al., 2017). However, the overall level of this motor-related activity is higher than one would expect simply from a motor-related prediction of visual input. Motor related activity is also evident in complete darkness and even prior to first visual experience in life (Attinger et al., 2017; Keller et al., 2012; Mahringer et al., 2022). It is possible, that this relatively high level of motor related activity in mouse visual cortex is a reflection of the fact that mice rely much less on fine visuomotor control.

In systems that have evolutionarily been driven to very high levels of precision of sensorimotor control, like vocal learning, there is much less such motor related activity. This is evident in the auditory system of a songbird, which relies on very temporally precise sensorimotor error detection for vocal learning, where there is much less motor related drive of auditory responses (Keller and Hahnloser, 2009). Similarly, there is very little movement related drive of activity in visual cortex of non-human primates (Liska et al., 2024; Talluri et al., 2023). In humans, movement-related modulation of EEG activity in visual cortex has been described (Cheng and Nordin, 2025; Gramann et al., 2010), but it is still unclear if visual responses are modulated by these signals. If indeed the evolutionary pressure for precision in visuomotor coupling determines the amount of motor modulation of visual responses, we would expect to find less of it in humans than in the mouse. Humans have a much higher reliance on visual feedback to guide movements than mice do.

This heightened reliance on visual guidance suggests that the human brain should be particularly adept at detecting mismatches in visuomotor coupling. Indeed, violations of the coupling between actual hand position and that of a virtual hand trigger responses in human visual cortex (Stanley and Miall, 2007). Even violations of simple action-outcome rules can trigger robust error responses in human subjects (Kimura and Takeda, 2019, 2014). Thus, we would also expect to find strong visuomotor mismatch responses in humans, comparable to – or stronger than – those observed in mice (Keller et al., 2012).

## RESULTS

### Visual responses

To quantify visual responses in freely moving human participants, we combined a virtual reality (VR) headset with an 8-electrode wireless EEG recording system (**Methods**; **Figure 1A**). Participants were shown a virtual environment that consisted of a large empty floor space with a square checkerboard pattern fixed to their viewing angle directly in front of them (**Figure 1B**). The checkerboard covered a visual angle of 53° and contained a red fixation dot in the center. Participants were asked to fixate on the red dot throughout all measurements. The checkerboard reversed white-black at intervals sampled from a uniform distribution between 2 and 4 s (**Figure 1C**). Recordings were split into a passive session during which participants were seated on a chair, and an active session during which they were instructed to walk around a rectangular floor area of 5 by 7 m. During the entire experiment, we tracked the location of participants (**Figure 1D**).

**Figure 1.**
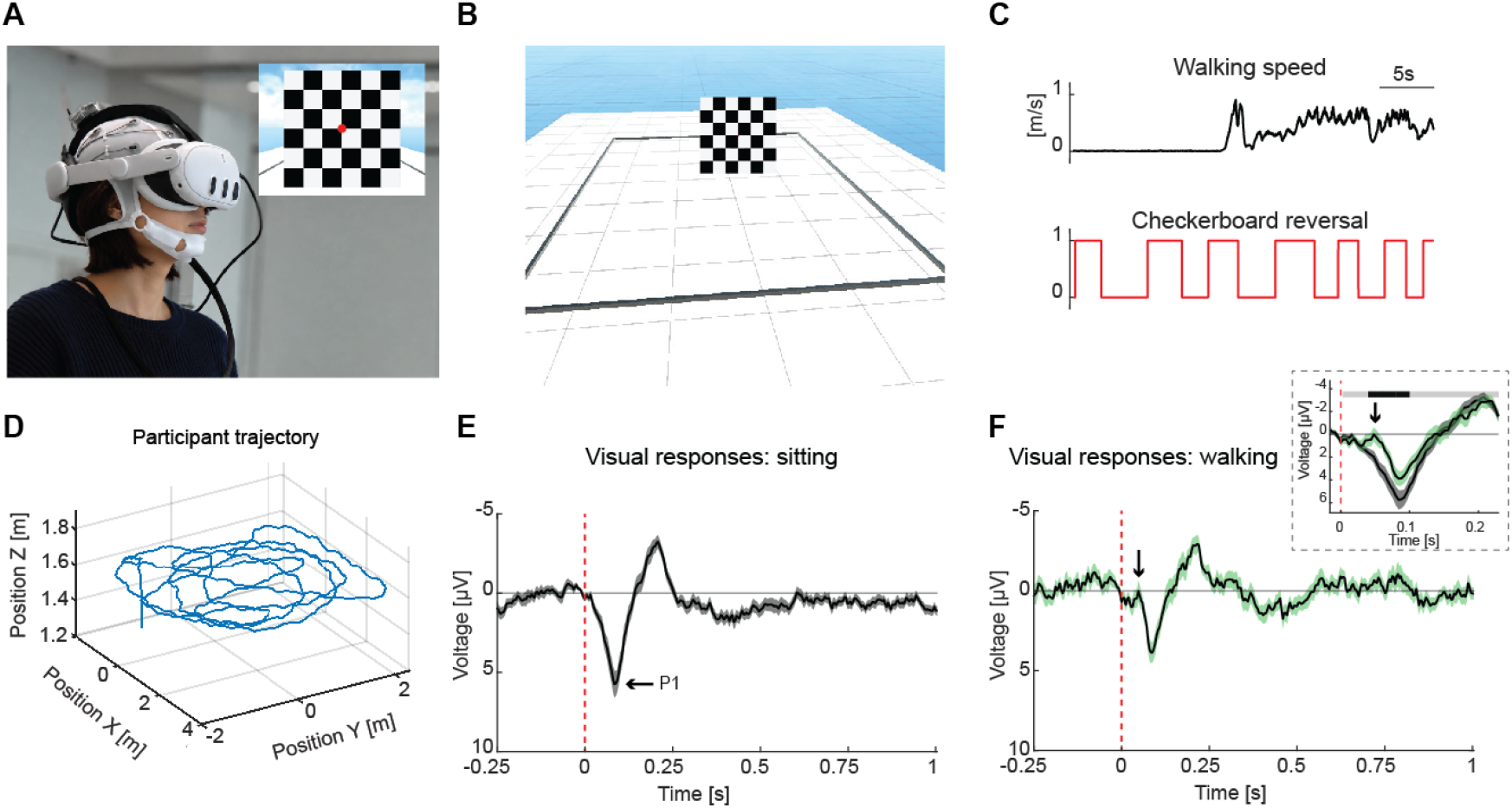
EEG responses to visual stimulation are modulated by walking. **(A)**Computer-generated image showing a participant wearing a wireless EEG recording setup and a virtual reality headset. An Open BCI EEG electrode cap is combined with META QUEST 3. Inset: First person view of the virtual environment. Red fixation dot was enlarged in figure to make it visible. **(B)**A third person view of the virtual scene used to study visual responses to a reversing checkerboard pattern. The black line on the ground indicates the safety boundary within which the participant was instructed to walk around during the movement block of the paradigm. The grid indicates 1 by 1 m squares. **(C)**Participants viewed a reversing checkerboard in two sessions: During half of the trials, they remained seated, and during the other half they walked within a defined safety boundary. Checkerboard reversed colors at random intervals every 2 to 4 s. **(D)**3D trajectory of an example participant during the visual experiment. Movement was recorded using accelerometers integrated into the VR headset. **(E)**Visually evoked potentials measured in the sitting session on the occipital electrodes O1 and O2. Responses recorded on other electrodes are presented in **Figure S3**. Solid black line represents the mean, and shading indicates the SEM across participants. Dashed vertical red line indicates the time of the checkerboard reversal. P1 - positive peak at around 88 ms. **(F)**As in **E**, but while participants were walking. Arrow indicates an early negative deflection preceding the P1 component. Inset: Comparison of visual evoked potentials measured in the sitting **E** and walking **F** sessions. The horizontal bar above the plot marks time bins in which responses differ significantly (black: p < 0.05) or do not differ (gray: p > 0.05).

Walking triggered movement related artifacts in the EEG signals (**Figure S1**). The strength of these movement artifacts varied as a function of hairstyle and gait of the participants. All participants were encouraged to go ‘gentle’ to reduce this problem. We excluded all data with eye blink and high movement related artifacts from further analysis (**Figure S2, Methods**; for number and percentage of excluded trials per session see **Table S2**).

We could detect multiphasic responses to the checkerboard inversion in occipital EEG electrodes (**Figures 1E and S3**). These responses were comparable to previous reports (Drislane, 2007) and exhibited a positive peak at around 88 ms (P1). Interestingly, when participants were walking, the same responses showed an early negative deflection with a peak latency of 48 ms that preceded P1 (**Figure 1F**). This early negative deflection at around 50 ms is also apparent in data from earlier studies comparing visual responses in walking and standing participants (Gramann et al., 2010).

While participants were instructed to fixate on the red dot, it is possible that there are systematic differences in eye movements between walking and sitting sessions. Our setup did not allow for concurrent eye tracking; thus, we cannot exclude the possibility that differences in eye movements contribute to the differences in EEG responses we observe. However, given the relatively rapid onset of response differences (40 ms), we suspect that stimulus-triggered eye movements cannot account for these early differences.

### Visuomotor mismatch responses

To measure visuomotor mismatch responses, participants were instructed to walk around in a virtual corridor (**Figure 2A**). The corridor was 1 m wide, 2.4 m high and of an oval shape covering approximately 7 by 5 m (**Figure 2B**). Virtual movement in the corridor was coupled to the movement of participants. We refer to this as closed loop coupling. To trigger visuomotor mismatches, we briefly (0.5 s) halted the coupling between movement of the participants and visual feedback in the virtual corridor at random times (every 10 to 15 s; **Methods**; **Figure 2C**). Throughout all experiments, we tracked the virtual position of participants as they were walking around the corridor (**Figure 2D**).

**Figure 2.**
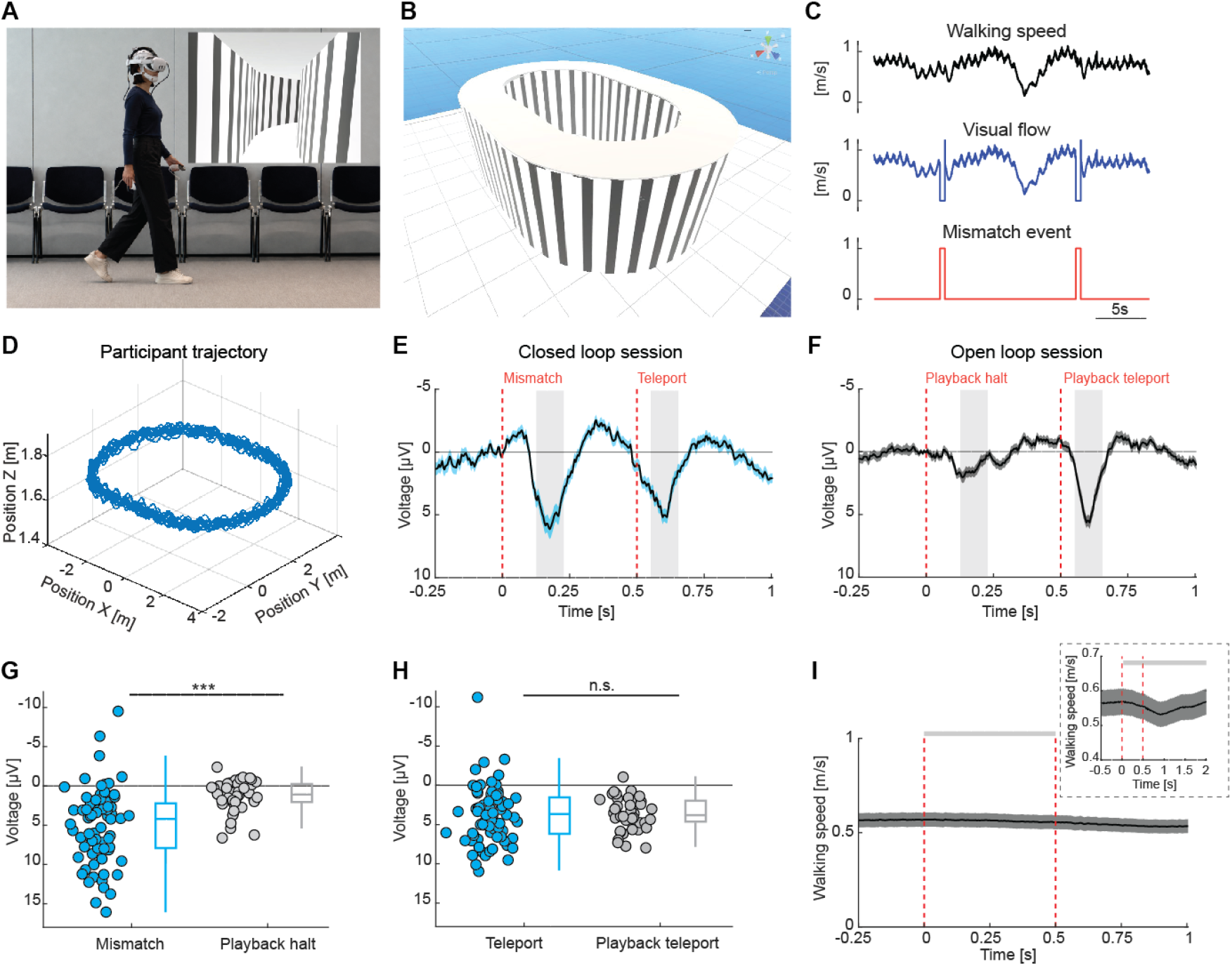
Visuomotor mismatch elicits strong EEG responses. **(A)**Computer-generated image showing a participant navigating a virtual tunnel. Inset: First person view of the tunnel. **(B)**Third person view of the virtual environment used to study visuomotor mismatch responses. The grid indicates 1 by 1 m squares. **(C)**In closed loop sessions, the walking speed of participants was coupled to movement in the virtual corridor. Visuomotor mismatches were introduced by briefly halting the visual flow for 0.5 s. Following each visuomotor mismatch event, the view was updated to the participants’ current position (brief peaks in visual flow speed after the mismatch event) and visuomotor coupling was resumed. **(D)**Trajectory of an example participant during the visuomotor mismatch paradigm. **(E)**Responses to visuomotor mismatches recorded from occipital electrodes (O1 and O2). Solid black line represents the mean, and shading indicates the SEM across participants. Dashed vertical red lines are onset and offset of the visuomotor mismatch. The gray shaded areas mark the analysis windows used to quantify response strength in **G** and **H**. Analysis windows of 100 ms were centered on the peak of the visuomotor mismatch response and the teleport response. **(F)**As in **E**, but for visual flow playback halt responses recorded from occipital electrodes. **(G)**Comparison of the response strength to visuomotor mismatches and playback halts. Here and elsewhere: Boxes mark median, quartiles, and range of data not considered outliers. Each circle represents data from one participant. ***: p<0.001. See **Table S1** for all statistical information. **(H)**Comparison of the response strength to teleport events following visuomotor mismatches in the closed loop session and playback halts in the open loop session. n.s.: not significant. See **Table S1** for all statistical information. **(I)**Average walking speed of participants during visuomotor mismatches. Inset: Temporally expanded view. The horizontal bar above the plot marks time bins where walking speed differs significantly (black: p < 0.05) or does not differ (gray: p > 0.05) from baseline.

Anecdotally, the perception some participants reported experiencing during these visuomotor mismatches was a very salient sudden movement of the entire visual world. In the words of one participant: “The world suddenly flew forward!”. This was reminiscent of a case report of a patient with a lesion to the lateral rectus muscle described by von Helmholtz (von Helmholtz, 1867): The lateral rectus muscle moves the eye laterally. The patient, upon attempting to move the affected eye laterally, reported seeing the world rapidly moving in the direction of the intended eye movement. One interpretation of this is that the combination of movement and absence of resulting visual flow change results in the perception that the world must be moving.

In the EEG recordings of occipital electrodes, we found strong responses triggered by these visuomotor mismatches (**Figure 2E**). Responses were dominated by a positive component peaking at 180 ms. Following the 0.5 s halting of the visual stimulus, the virtual location was updated to match the actual location of participants (we call this a teleport event), and the visual flow coupling was resumed. This resulted in a sudden change of visual stimulus combined with a visual flow onset and drove what is likely a visual response in the EEG signal (**Figure 2E**).

To quantify how much of the visuomotor mismatch response can be explained by visual input alone, we exposed participants to a replay of the visual flow that was self-generated in the preceding closed loop session, including the visual flow halts that constitute visuomotor mismatches in the closed loop session. We refer to this as open loop coupling. For these experiments, participants were seated on a chair. To reduce the likelihood of triggering nausea in participants, roll and pitch movements of the head were removed from this replay. We found measurable responses to visual flow playback halts (**Figure 2F**), but these were much smaller than visuomotor mismatch responses (**Figure 2G**). In the few participants who volunteered to experience playback that included pitch and roll movements, playback halt responses were also significantly smaller than visuomotor mismatch responses (**Figure S4A-S4C**) and not different from those in the reduced open loop session (**Figure S4D and S4E**). The difference in response amplitude between mismatch and playback response could not be explained by a simple movement related gain of visual responses, as the response to the teleport and re-onset of visual flow was similar in both closed and open loop sessions (**Figure 2H**). Note that the mismatch responses and playback halt responses shown here do not come from fully overlapping pools of participants (**Table S1**). A subset of visuomotor mismatch sessions had to be excluded because of recording noise, while some participants aborted the open loop recording due to developing nausea (**Methods**; for number and percentage of excluded trials per session see **Table S2**). A participant-wise comparison of mismatch and playback halt responses is shown in **Figure S5**.

By design, open loop sessions in the form of an identical playback (to match the visual statistics of open and closed loop sessions) always followed closed loop sessions, which could mean that the difference between visuomotor mismatch and playback halt responses could be driven by some form of adaptation. To control for this, we repeated the experiment in a separate cohort of participants with the order of sessions reversed: Participants first viewed a playback of the closed loop session of the previous participant and only then performed an active closed loop session. We found that independent of the order of closed and open loop sessions, visuomotor mismatch responses were larger than playback halt responses (**Figure S6**), thus excluding the possibility that the response difference may be driven by adaption or a reduction in salience of the sudden flow halt.

Although participants reduced their walking speed in response to the visuomotor mismatch when averaged over all trials, this did not reach statistical significance (**Figure 2I**). But see **Figure 4A**, for evidence that there was a significant reduction in walking speed when analyzing only the first few mismatch trials. In either case however, the reduction in walking speed was of much longer latency compared to the visuomotor mismatch responses, demonstrating that walking speed changes cannot explain visuomotor mismatch responses.

To test whether visuomotor mismatch events trigger eye movements that could explain visuomotor mismatch responses, we measured eye movements triggered by visuomotor mismatch in an independent cohort of participants. As with the EEG experiments, participants were exposed to a closed loop session with visuomotor mismatches. Throughout the experiment, we tracked eye position and eye blinks using an eye tracker built into the VR headset (**Methods**). Note, this VR headset was different from the one we used in the EEG experiments. We found that visuomotor mismatch triggered detectable eye responses. While we could not detect any systematic eye movements triggered by mismatch (**Figure S7A and S7B**), participants exhibited a reduction in average eye velocity (**Figure S7C**) and a reduction in eye blink rate (**Figure S7D**) with a latency of about 208 ms and 541 ms, respectively. Thus, both of these changes had latencies longer than those we observed for responses to visuomotor mismatch in the EEG experiments (**Figure S7E**). Thus, we find no evidence that visuomotor mismatch responses can be explained by eye blinks or eye movements.

Finally, we performed time-frequency power and phase-locking analyses of visuomotor mismatch and playback halt responses. The time-frequency power analysis revealed an increase in delta-band power during visuomotor mismatch, consistent with previous reports linking delta activity to prediction error processing, including reward prediction errors (Cavanagh, 2015), unexpected final words (Webb and Sohoglu, 2025), and visual deviance detection (West et al., 2024). Notably, it appears as if the increase in delta power emerged first over occipital electrodes and appeared later over more frontal electrodes, forming a spatiotemporal gradient of onset across the scalp (**Figure S8**). Delta power changes were markedly reduced in the playback halt responses and power changes occurred primarily at visual flow onset rather than at flow offset. The inter-trial phase coherence analysis revealed a strong delta-band synchronization over occipital electrodes following visuomotor mismatch, while the playback halt response primarily resulted in phase synchronization in both delta and theta bands following visual flow onset (**Figure S9**). These findings suggest that visuomotor mismatch evokes a delta-dominated low-frequency response that emerges first over occipital electrodes, then spreads across the cortex, and cannot be explained by passive visual flow changes alone. Unlike auditory oddball MMN responses, which show transient increases across a broader 3-12 Hz range (Ko et al., 2012), our response is more selectively delta-band dominant. This pattern more closely resembles visual deviance responses (West et al., 2024), where delta activity was stronger than theta and more prolonged, matching the temporal dynamics observed in our data.

### Distribution of visuomotor mismatch response strength

In mice, visuomotor mismatch responses originate in primary visual cortex and propagate to a network of areas across dorsal cortex (Heindorf and Keller, 2023; Takeuchi et al., 2024). Thus, we expected to find larger and faster responses in the two occipital electrodes. To test this, we compared the visuomotor mismatch responses across the eight recorded locations in both response strength and timing (**Figures 3A and 3B**). We indeed found that the strongest responses were observed at occipital electrodes O1 and O2, though significant responses were also present at frontal (Fp1-2) and parietal (P3-4) sites (**Figure 3C**). We found a significant difference in response latency between occipital and frontal electrodes, with occipital signals occurring significantly earlier than frontal responses. (**Figures 3D and 3E**). This suggests that mismatch processing may initially arise in sensory visual areas before engaging higher-order frontal regions.

**Figure 3.**
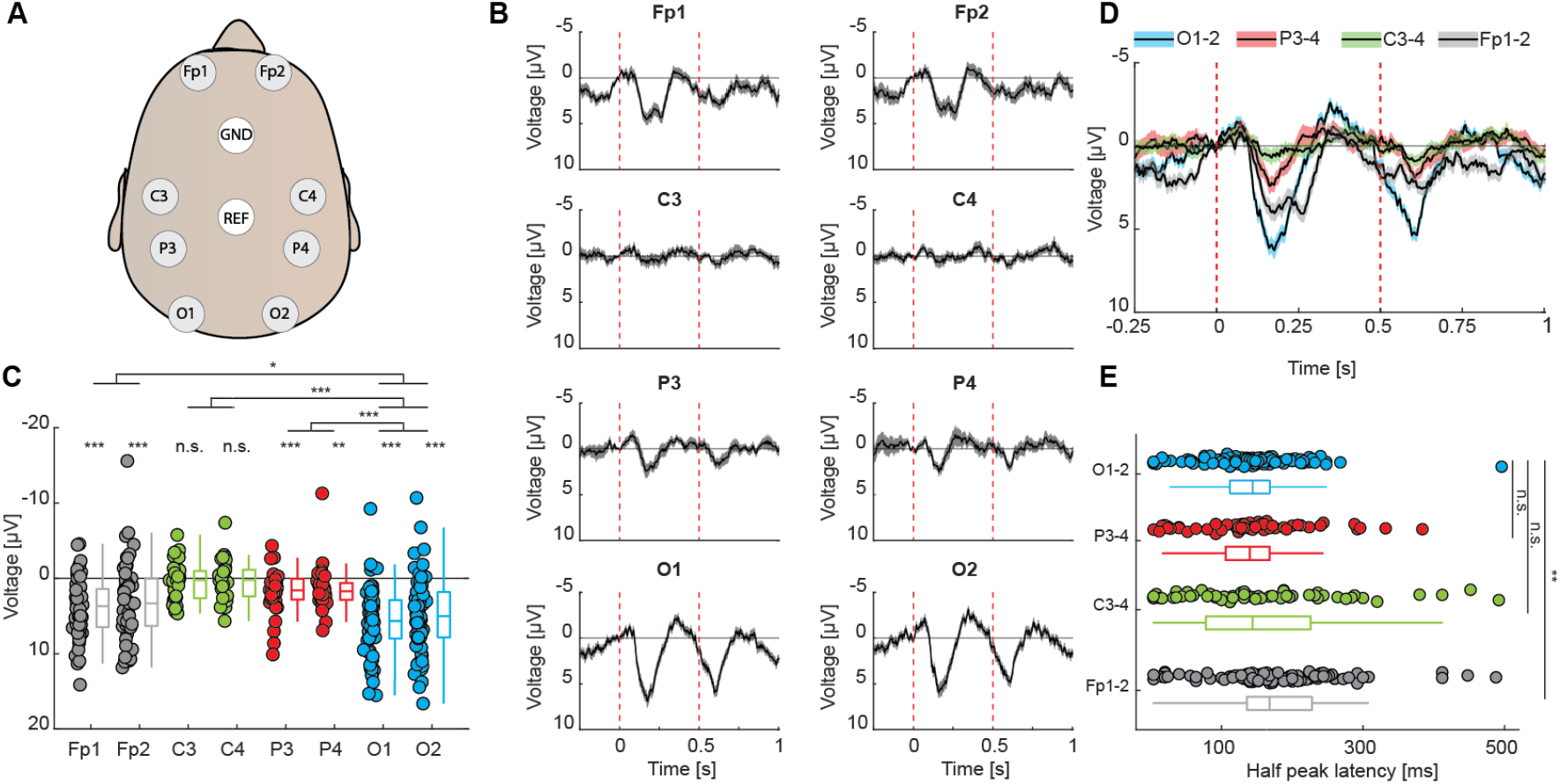
Visuomotor mismatch responses are most prominent over occipital cortex. **(A)**Top down view of EEG electrode locations on the head. **(B)**Visuomotor mismatch responses measured on electrodes shown in **A**. Solid black lines represent the mean, and shading indicates the SEM across participants. Dashed vertical red lines are onset and offset of the visuomotor mismatch. **(C)**Comparison of the response strength to visuomotor mismatches measured on electrodes shown in **A**. Average response strength was calculated within a 100 ms window centered on the peak of the average visuomotor mismatch response across all electrodes. Boxes mark median, quartiles, and range of data not considered outliers. Each circle represents data from an individual participant. ***: p<0.001, **: p<0.01, *: p<0.05, n.s.: not significant. See **Table S1** for all statistical information. **(D)**The responses shown in **B**, averaged over pairs of electrodes and overlaid. Solid black lines represent mean and shading SEM across electrodes. Dashed vertical red lines are onset and offset of the visuomotor mismatch. **(E)**Comparison of the latency to half maximum response for the 4 electrode pairs. **: p<0.01, n.s.: not significant. See **Table S1** for all statistical information.

### Temporal dynamics of neuronal and behavioral responses to visuomotor mismatch across a session

To examine how mismatch responses evolved over the session, we compared responses in early (first five) and late (last five) trials. We found that participants reduced their walking speed with a delay of about 500 ms following mismatch events. This effect was more prominent in early trials (14.3%) compared to late trials (5.7%) (**Figure 4A**). Interestingly, visuomotor mismatch responses mirrored this pattern over frontal electrodes with a clear attenuation from early to late trials (**Figure 4B**). By contrast, response changes in occipital electrodes were smaller between early and late trials (**Figure 4C**). The fact that the reduction over frontal cortex is larger than that of responses over visual cortex (**Figure 4D**), indicates that there may be a gradient of experience dependent changes in processing from early sensory to frontal areas of cortex that relate to changes in behavioral responses to, but not detection of prediction errors.

**Figure 4.**
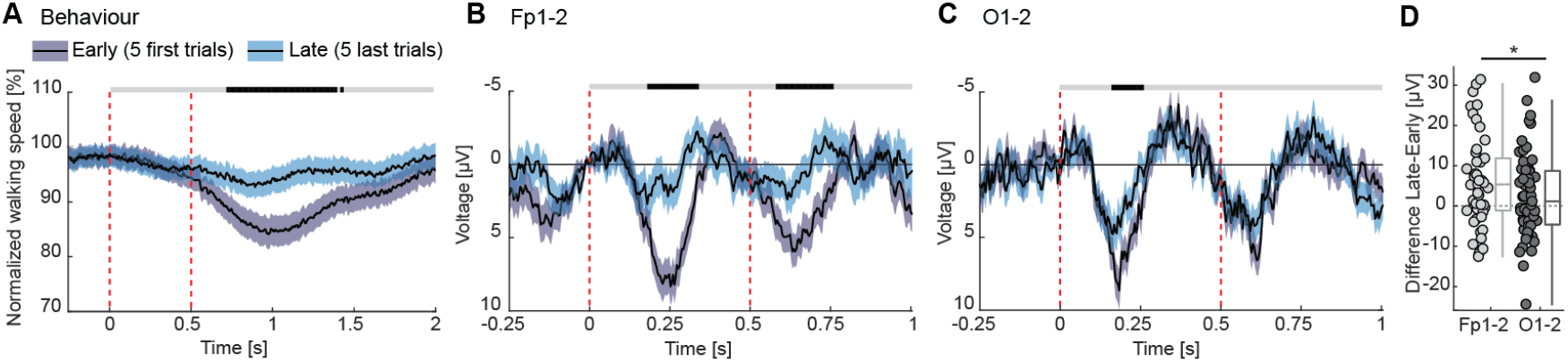
Visuomotor mismatch responses show experience-related changes predominantly over frontal electrodes. **(A)**Average walking speed during visuomotor mismatch events in early versus late trials. The solid black line shows the mean and the shaded area the SEM across participants. Vertical red dashed lines indicate mismatch onset and offset. The horizontal bar above the plot marks time bins in which the walking speed in early trials differs significantly (black, p < 0.05) or not (gray, p > 0.05) from that in late trials. **(B)**Visuomotor mismatch responses in early versus late trials over frontal electrodes. **(C)**As in **B**, but for occipital electrodes. **(D)**Comparison of the difference between visuomotor mismatch response in early and late trials for frontal and occipital electrodes. Average response strength was calculated within a 100 ms window centered on the peak of the average visuomotor mismatch response in early trials above frontal or occipital electrodes. Boxes mark median, quartiles, and range of data not considered outliers. *: p<0.05. See **Table S1** for all statistical information.

### Comparison of visual responses and visuomotor mismatch responses

The time course of visual responses (**Figure 1E**) and visuomotor mismatch responses (**Figure 2E**) differed in that they appeared to exhibit reversed polarity (**Figure 5A**). Despite this difference in polarity, the initial peaks occurred at similar latencies. The total power of the visuomotor response was higher than that of the visual response (**Figure 5B**), but given that we did not optimize either the visuomotor mismatch stimulus nor the visual stimulus for maximum response, this comparison is more qualitative than quantitative in nature. We include it here primarily to illustrate how the visuomotor mismatch response compares to a more conventional visual response.

**Figure 5.**
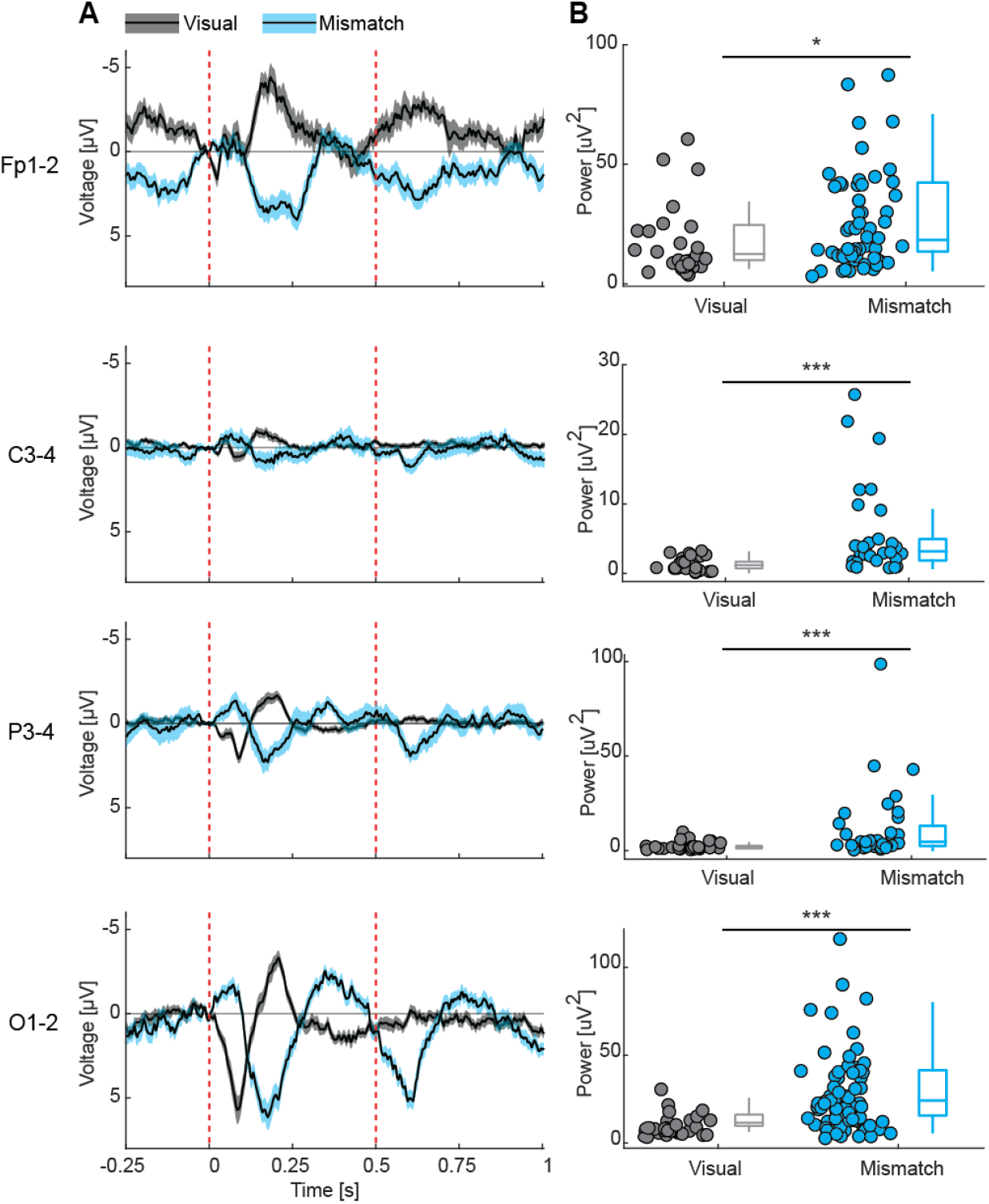
Visuomotor mismatch responses have reversed polarity and more power compared to visual responses. **(A)**Comparison of visual (**Figure 1E**) and visuomotor mismatch (**Figure 2E**) responses recorded from all electrodes. Solid black line represents the mean, and shading indicates the SEM across participants. Dashed vertical red lines are onset (visual and mismatch) and offset (mismatch) of the stimuli. **(B)**Comparison of the power of visual and visuomotor mismatch responses, calculated within a 0 - 0.5 s time window following stimulus onset. Boxes mark median, quartiles, and range of data not considered outliers. Each circle represents data from an individual participant. ***: p<0.001, *: p<0.05. See **Table S1** for all statistical information.

### Comparison of visuomotor mismatch responses and auditory oddball mismatch responses

To further contrast the strength of visuomotor mismatch to other frequently used prediction error signals, we also recorded EEG responses in an auditory oddball paradigm. Mismatch responses recorded with this paradigm can be thought of as a stimulus history prediction error (Garrido et al., 2009; Näätänen et al., 2007). We implemented an oddball paradigm composed of a sequence of identical tones that was occasionally disrupted by a tone of a different frequency (**Figure 6A**). We used pure tones of 1 kHz and 1.2 kHz frequency and alternated in blocks which tone was standard and which was the deviant (**Methods**). Participants watched short silent movies in the VR headset while listening to the tone sequences delivered in headphones. To isolate the oddball driven mismatch response, we subtracted the response to the tone when presented as a standard from the response to the same tone when presented as a deviant. The resulting difference waveform closely resembled those reported in previous studies (**Figures 6B-6C, S10, and S11**) (Näätänen et al., 2007). Comparing these oddball mismatch responses to the visuomotor mismatch response in occipital electrodes, we found that both had similar dynamics with a dominant positive peak at around 180 ms (**Figure 6D**). Comparing the total power of the two responses, we found that visuomotor mismatch responses were significantly larger than oddball mismatch responses (**Figure 6E**). This was also true for responses measured in temporal electrodes (**Figures S10 and S11**). But also here, the comparison of total power is more qualitative than quantitative as neither stimulus was optimized for maximum EEG response, and we include it to illustrate how typical variants of these responses compare.

**Figure 6.**
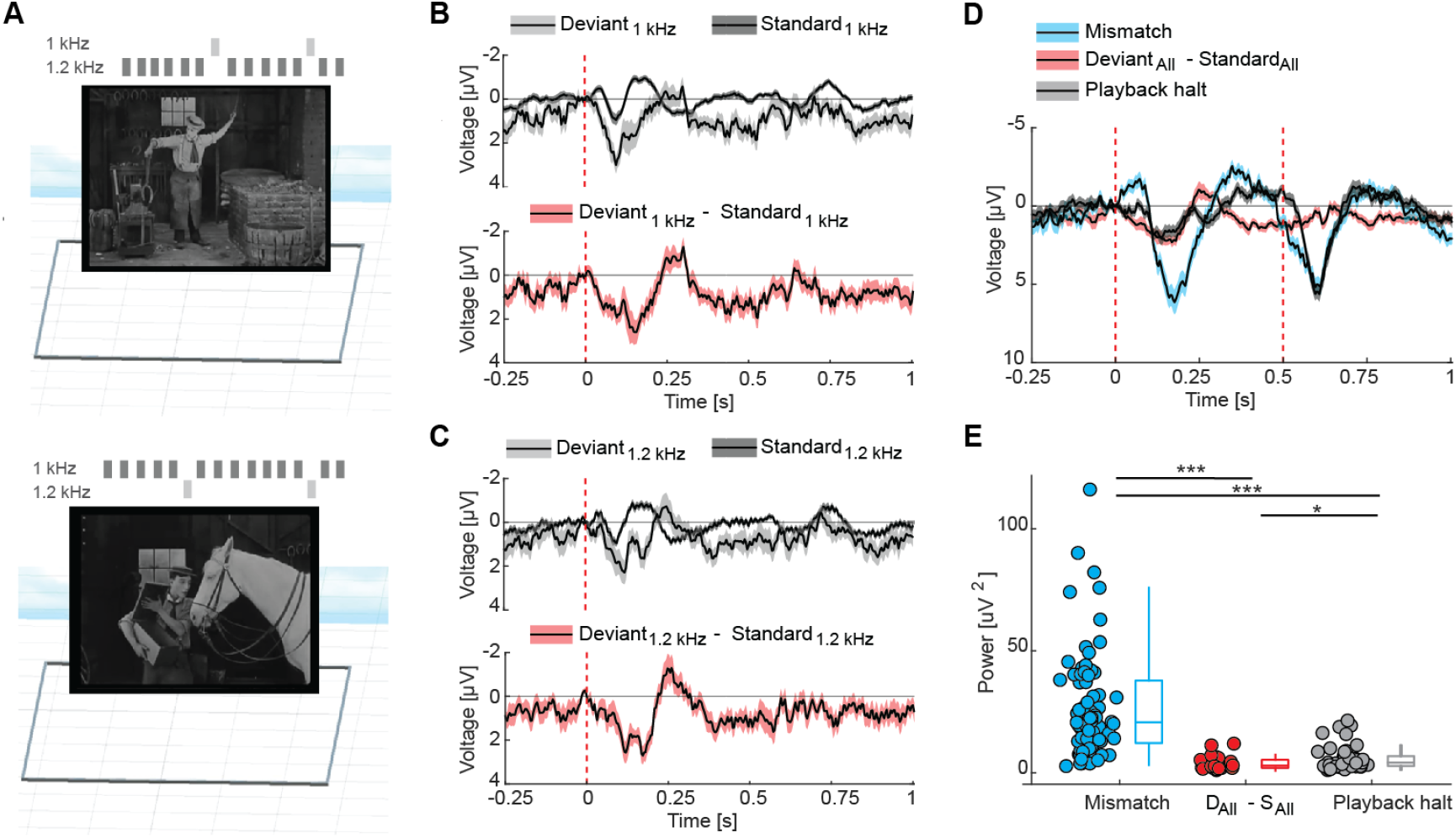
Visuomotor mismatch responses are larger than auditory oddball mismatch responses but have similar temporal dynamics. **(A)**Design of the auditory oddball paradigm with examples of the silent films participants were exposed to. **(B)**Top: Auditory responses to the 1 kHz tone presented as a standard versus as a deviant. Bottom: Oddball mismatch response calculated by subtracting the average response over trials in which the tone was presented as a standard from those when the tone was presented as a deviant. Solid black lines represent the mean, and shading indicates the SEM across participants. Dashed vertical red line is the onset of the auditory stimulus. **(C)**As in **B**, but for the 1.2 kHz tone. **(D)**Comparison of visuomotor mismatch, oddball mismatch (average over data shown in panels B and C) and playback halt responses recorded from occipital electrodes. Solid black lines represent the mean response, with shading indicating SEM across electrodes. Dashed vertical red lines are onset (visual, mismatch) and offset (mismatch) of the stimuli. **(E)**Comparison of the power of visuomotor mismatch, oddball mismatch response (average over data shown in panels B and C) and playback halt responses, calculated within a 0 s - 0.5 s time window following stimulus onset. Boxes mark median, quartiles, and range of data not considered outliers. Each circle represents data from an individual participant. ***: p<0.001; *p<0.05. See **Table S1** for all statistical information.

## DISCUSSION

The utility of paradigms for measuring prediction errors in humans primarily comes from their relevance to psychiatric disorders, where disruptions in prediction error processing are widely implicated (Adams et al., 2013; Fletcher and Frith, 2009; Kirihara et al., 2020; Qela et al., 2025; Sterzer et al., 2018). Various types of prediction error responses are being explored as potential biomarkers in clinical research, serving to monitor disease progression, evaluate symptoms, and provide insight into the underlying neuronal mechanisms of these conditions.

The most prominent of these prediction errors is mismatch negativity (MMN). MMN is the first component of the oddball mismatch response (Näätänen et al., 1978). MMN has been shown to be reduced or abnormal in patients with schizophrenia spectrum disorders (Todd et al., 2012; Umbricht and Krljes, 2005), autism spectrum disorders (Chen et al., 2020; Dunn et al., 2008; Schwartz et al., 2018), speech disorders (El Hatal de Souza et al., 2020), depression (Tseng et al., 2021), and bipolar disorders (Chitty et al., 2013). More generally, it is thought that many of these disorders can be related to changes in predictive processing. In the case of schizophrenia, for example, the symptoms are thought to arise from an imbalance in the strength of high and low level predictions (priors) (Schmack et al., 2017, 2015a, 2015b, 2013). Further testing these ideas will require experiments that trigger identified functional responses in humans, for which we have a circuit level understanding based on cell type specific recordings.

We were able to measure robust visuomotor mismatch responses in freely moving humans (**Figure 2**). These signals cannot be attributed to changes in participants’ behavior or to simple visual offset responses. Notably, the visuomotor mismatch responses exhibited a markedly different temporal profile compared to purely visual responses (**Figure 5**). What might account for this difference? One possibility is that visuomotor mismatch signals may rely more strongly on input from other cortical areas than visual responses and are thus delayed relative to visual responses. A more interesting interpretation is that visual responses and visuomotor mismatch responses are both prediction errors of different types. A central tenet of the cortical circuit for predictive processing is the split into separate populations of neurons that compute positive and negative prediction errors (Keller and Mrsic-Flogel, 2018; Rao and Ballard, 1999). In this interpretation, a visuomotor mismatch response is a negative prediction error, while the response to a visual stimulus is a positive prediction error.

Visuomotor prediction errors are likely computed in layer 2/3 of mouse visual cortex (Jordan and Keller, 2020), and are also preferentially detectable in superficial layers of human visual cortex (Thomas et al., 2024). In the mouse, positive and negative prediction errors are likely computed in separate populations of neurons in visual cortex (Keller and Mrsic-Flogel, 2018; O’Toole et al., 2023). These two cell types have a different depth distribution, with negative prediction error neurons more superficial and positive prediction error neurons located deeper in cortex (O’Toole et al., 2023). Given that EEG signals are thought to exhibit positive or negative deflections as a function of depth of the source (Cohen, 2017; Kirschstein and Köhling, 2009), the polarity reversal of visual and visuomotor mismatch responses (**Figure 5**) may be the consequence of different populations of cells being activated and inhibited. Visuomotor mismatch should activate negative prediction error neurons and inhibit positive prediction error neurons, while the reverse is the case for a visual stimulus.

The fact that there is also a strong visual response to the visual flow re-onset following visuomotor mismatch means that our visuomotor mismatch paradigm might allow us to measure both negative and positive prediction error responses. More intriguingly, there is a differential modulation by walking of the two responses (**Figures 2G and 2H**). In theory, visuomotor mismatch drives a combination of two negative prediction errors. One is based on a movement related prediction. The other on a stimulus-history related prediction. Both the self-motion as well as the ongoing visual flow would predict a continuation of visual flow. A visuomotor mismatch violates both predictions. During passive observation, there is no movement-related prediction, but the stimulus history-based prediction is still violated. This could explain the strong reduction of the negative prediction error response during sitting, and may also explain why there is no reduction for the positive prediction error response.

Visuomotor mismatch responses recorded over frontal electrodes adapted more strongly with experience than responses recorded over occipital electrodes (**Figure 4**). This may suggest that sensory areas primarily act as detectors of visuomotor mismatch events (subjective prediction error signal), whereas frontal areas reflect higher-level adaptation to the event and the resulting behavioral adjustment. Mouse data suggested that visuomotor mismatch responses originate in visual sensory areas (Heindorf and Keller, 2023; Jordan and Keller, 2020) and are broadcast to other cortical areas from there. Thus, the stronger decrease over frontal cortex suggests that signals measured over frontal electrodes may reflect the integration of sensory mismatch information with behavioral or attentional signals. This behavioral component makes the paradigm particularly interesting as a potential clinical biomarker, as it may allow us to assess how quickly participants adapt their behavior to mismatch event and how this adaptation relates to prediction error responses in frontal cortex.

Intriguing is the similarity in timing and polarity of the playback halt and oddball response (**Figure 6D**). An oddball response is always a combination of both a positive and a negative prediction error. There is a negative prediction error for the absence of the expected standard tone as well as a positive prediction error for the presence of the unexpected oddball tone. Thus, it is conceivable that the subtraction of standard response from the oddball response, isolates the component of the oddball response that corresponds to a negative prediction error.

One of the key challenges in systems neuroscience is translating findings from animal models to humans. Although recent animal studies have provided detailed insights into the circuit level implementation of predictive processing in the cortex (Keller and Mrsic-Flogel, 2018), and psychological and psychiatric conditions have long been described within this framework (Keller and Sterzer, 2024; Sterzer et al., 2018), translation to human research has been limited by a lack of methods to record cell type specific signals in human experiments. With an understanding from animal models of how positive and negative prediction errors are computed, and which cortical layers are involved, an approach based on functionally identified responses might be most promising. One step in this direction is to use experimental paradigms that can cleanly separate e.g. positive and negative prediction error responses. As argued above, we speculate that the visuomotor mismatch paradigm as we have used it here, is one possible way to achieve this.

It should also be highlighted that the paradigm we have developed here is still limited in its clinical applicability by practical constraints, including movement-related artifacts, signal-to-noise levels, low channel density, and the requirement for active engagement. Nevertheless, active task engagement may itself be a critical feature for revealing deficits in cortical processing that remain undetectable in more passive paradigms. Thus, although active, more video-game-like paradigms may appear less practical in certain clinical contexts, they may be essential for capturing meaningful and behaviorally relevant neuronal responses. It will also be of importance to overcome a challenge of performing recordings with a higher channel density in freely moving participants for improved source localization. Thus, methodological refinement and hardware optimization will be required before reliable clinical translation. However, these limitations do not detract from the paradigm’s immediate utility as a research tool for studying prediction error processing.

## METHODS

### Participants

The study was approved by Ethikkommission Nordwest-und Zentralschweiz (Project-ID: 2024-02458). Participants signed a written informed consent before participation and were not financially compensated for their participation. A total of 91 recording sessions from healthy adults were included in the study. Sessions were obtained from participants aged 18–65 years, with a small number of participants contributing more than one recording session. The age distribution across sessions was as follows: 38 sessions from ages 18–30, 37 sessions from ages 31–40, 11 sessions from ages 41–50, and 5 sessions from ages 51–65. None of the participants reported a prior diagnosis of movement disorders, vestibular dysfunction, or epilepsy. Experience with virtual reality technology ranged from beginner to advanced, with most participants reporting minimal prior experience.

### EEG recordings in humans

We integrated a wireless EEG recording system with a VR headset. The EEG recording system was composed of a wet electrode cap and a Cyton biosensing board from OpenBCI. This allowed us to record 8 EEG channels (43 sessions: Fp1, Fp2, C3, C4, P3, P4, O1 and O2; 48 sessions: Fp1, Fp2, T3, T4, T5, T6, O1 and O2) at a 250 Hz sampling rate. Electrode labels follow the international 10-20 system: Fp = frontopolar, C = central, P = parietal, O = occipital, and T = temporal; odd numbers indicate the left hemisphere, and even numbers the right (**Figure 3A**). All EEG data were wirelessly (via Bluetooth) transmitted to a nearby computer. The VR headset was a Meta Quest 3, with a virtual environment developed in the Unity engine (Unity Technologies). Virtual 3D objects were designed in Fusion 360 (Autodesk). Synchronization between EEG recording and VR headset was performed by connecting an auditory output of the VR headset to the Cyton board to exchange synchronization triggers. This allowed us to synchronize EEG data with VR events offline. Auxiliary signals, including the participant’s position, stimulus trigger timing, and type, were recorded directly on the headset at a sampling rate of approximately 100 Hz.

### Eye movement measurements

We recorded eye blinks and eye movements using a Meta Quest Pro VR headset and its built in eye tracker during closed loop sessions. Eye-tracking data were sampled at 72 Hz. We combined blink signals from left and right eye, by averaging the two channels at each time point, yielding a single blink/eye-closure trace per participant. To compute eye movement speed, we first computed eye-movement speed separately for the left and right eye from the moment-to-moment change in eye rotation between consecutive samples and then averaged the two speed traces to obtain a single eye-speed measure.

### Visual responses

Visual stimuli were presented using the VR headset. We used a reversing square checkerboard stimulus to drive visually evoked potentials. The checkerboard reversed colors at random intervals (between 2 and 4 s). In virtual space, the checkerboard measured 0.5 by 0.5 m and was positioned 0.5 m in front of participants’ eyes. This resulted in a horizontal and vertical coverage by the checkerboard of approximately 53° of visual angle. Each visual stimulation session lasted 4 min, during which the checkerboard reversed colors between 78 and 82 times. For half of the session, participants viewed the stimulus while seated; for the other half, they were instructed to walk freely within a 7 by 5 m empty floor space. The order of sitting and walking sessions was randomized across participants.

### Visuomotor mismatch responses

For the experiments measuring visuomotor mismatch responses, we used a 3D virtual corridor with vertical gratings on the walls. The corridor measured 1 m in width, 2.4 m in height, and had an oval shape of 7 by 5 m. Visuomotor mismatches were introduced at random intervals every 10 to 15 s as participants walked through the corridor. The session lasted 5 min and included between 22 and 26 visuomotor mismatch events. During these events, the coupling between the participant’s movement and the visual feedback in the VR headset was briefly interrupted, i.e. the visual scene was frozen for 0.5 s. Because participants continued to move during this time, they were teleported to their current position in the virtual space following visuomotor mismatch event.

### Playback halt responses

To quantify how much of the visuomotor mismatch response could be explained by visual input alone, participants were asked to passively observe a replay of the visual flow they had self-generated during the preceding closed loop session; we refer to this replay condition as the open loop session. We quickly learned, however, that watching 5 minutes of playback in the VR headset triggered nausea in most participants. Thus, we started experimenting with changes to the playback to minimize the risk of triggering nausea. One such modification we settled on to use for experiments was a playback version that omitted pitch and roll movements of the head. Full playback involved six degrees of freedom (6DOF): 3D position in space, plus pitch, yaw, and roll angles of the head. The constrained playback included only four degrees of freedom (4DOF): The 3D position in space, but only yaw movements of the head. A subset of participants volunteered to view the full playback (**Figure S4**). Open loop sessions were conducted while participants were seated and lasted 5 min, matching the duration of the closed loop session. Approximately 23% of participants reported strong nausea during the beginning of the 4DOF open loop session, at which point the recordings were terminated.

### Mismatch negativity

To measure mismatch negativity, we used an auditory oddball paradigm. Auditory stimuli consisted of two pure tones: 1 kHz and 1.2 kHz. They were presented at 60 to 70 dB sound pressure level in a counterbalanced design, in which each tone served as the standard in one condition and as the deviant in the other. Tones were 50 ms in duration, including 5 ms linear rise and fall ramps. Stimuli were delivered in a pseudorandom order, with deviant tones comprising 10% of all trials and preceded by at least five standard tones. The inter-trial interval was selected from a uniform distribution of between 500 and 600 ms. Each session lasted 3 min and included between 28 - 33 deviant and 244 - 299 standard stimulus presentations. Participants were seated and listened to the tones while watching a silent movie that was not related to the tone sequences.

### EEG Signal Analysis

Data analysis was done using custom-written MATLAB scripts. EEG signals were band-pass filtered between 1 and 100 Hz. To remove power line noise, a band-stop filter was applied between 40 and 60 Hz. Movement of the participants triggered all varieties of movement related artifacts in the EEG recordings. The strength of these artifacts depended on a variety of factors: Impedance of the electrodes, hair style of participants, gait pattern, and likely others. To reduce data contaminated by excessive movement artifacts and eye blinks, trials with a maximum absolute response amplitude exceeding 100 µV within the time window −0.5 to 1 s relative to the trigger were discarded from further analysis (**Figure S2**). Data from each electrode were included in the final analysis only if at least 15 triggers remained after exclusion of triggers with excessive movement artifacts (**Table S1**). To compare average response strength across sessions, a 100 ms analysis window was used, centered on the peak of the respective responses: The visuomotor mismatch event recorded at occipital electrodes O1 and O2 (**Figures 2G, S4C, S5C, S6C**), the visual flow re-onset response (**Figure 2H**), the visuomotor mismatch response in early trials for frontal and occipital electrodes (**Figure 4D**), the mean visuomotor mismatch response across all electrodes (**Figure 3C**), the mean visual response across all electrodes (**Figure S3C**) or playback halt response in the 6DOF session (**Figure S4E**). Signal power was compared by calculating the mean squared amplitude within a 0 - 0.5 s analysis window following stimulus onset (**Figures 5B, 6E, S10E, S11D**). To analyze how responses evolved over the course of the session, we computed moving-average responses across 10 trials at successive percentages of progress through the paradigm (**Figure S6D and S6E**). Time–frequency power and inter-trial phase coherence (ITPC) were computed using complex Morlet wavelet convolution approach (Cohen, 2014). EEG data were sampled at 250 Hz and analyzed from −0.5 to 2 s relative to the event of interest. Wavelets were generated from −1 to 1 s and covered 30 logarithmically spaced frequencies between 2 and 40 Hz. The number of cycles increased logarithmically with frequency, from 3 to 10 cycles.

### Statistical tests

All statistical analyses were conducted using hierarchical bootstrap (Saravanan et al., 2020). Bootstrap resampling enables statistical comparisons across sessions without assuming a specific distribution of the EEG data. For analysis in **Figures 1, 2, 4, 6, S4, S5 and S6**, we averaged signals across electrode pairs (O1-2 or Fp1-2) and treated the result as a single data point per participant. For **Figures S10 and S11**, the same procedure was applied to temporal electrode pairs (T3-4 and T5-6). Likewise, in **Figures 3D, 3E, and 5**, we averaged signals from the respective electrode pairs O1-2, C3-4, P3-4, and Fp1-2, and treated each pair as a single data point per participant. For the analysis shown in **Figures 3C and S3C** we included signals from all electrodes and used a nested bootstrap to account for multiple data points originating from the same participant. We first resampled the data with replacement at the level of participants, followed by resampling at the level of electrodes. For each resampled population, we computed the mean response and repeated this procedure 10000 times. The p value was estimated as the fraction of bootstrap samples in which the sample mean violated the tested hypothesis. See **Table S1** for all information on number of participants or electrodes used for all analyses shown.

## ACKNOWLEDGEMENTS

We thank all participants for taking part in the study. We thank all members of the Keller lab for discussion and support. We thank Philipp Sterzer for feedback on the manuscript. This project has received funding from the Swiss National Science Foundation (GBK), the Novartis Research Foundation (GBK), and the European Research Council (ERC) under the European Union’s Horizon 2020 research and innovation programme (grant agreement No 865617) (GBK).

---

**Figure S1.**
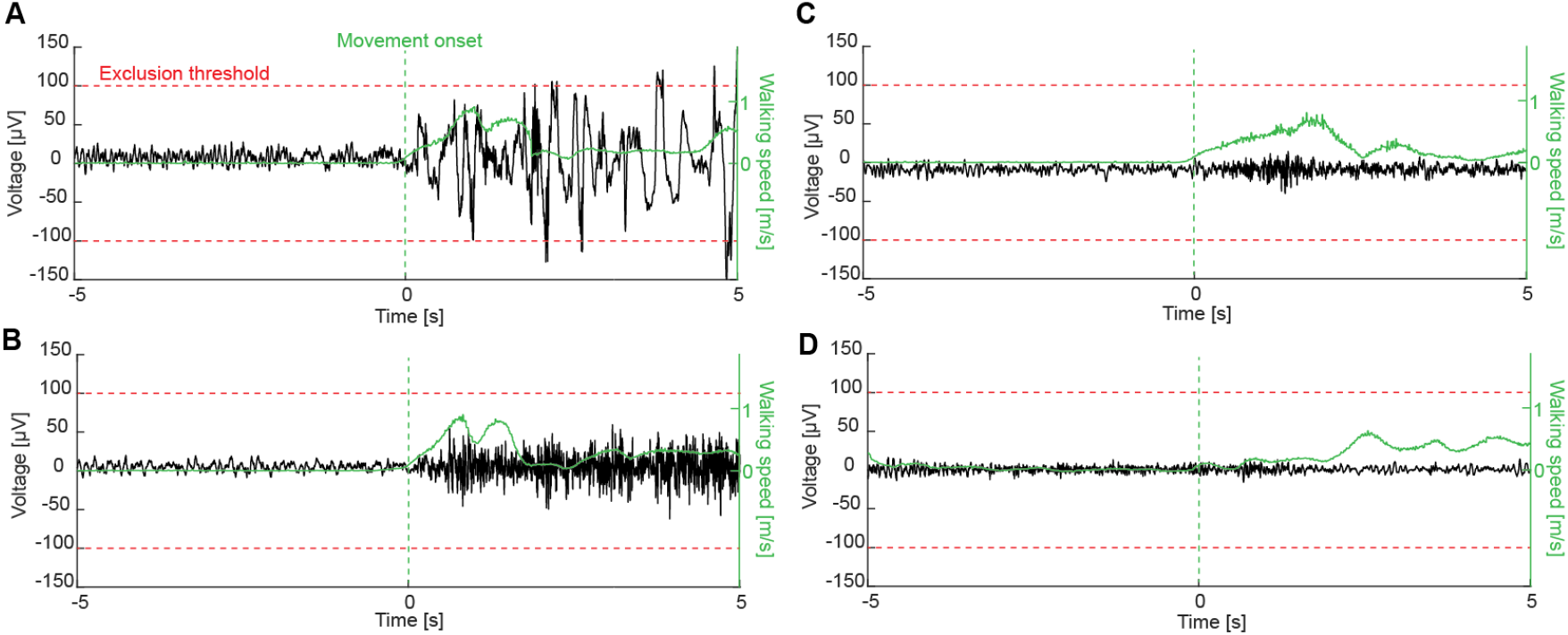
Movement onsets result in increases in variance in EEG activity. **(A)**We included only data in which the EEG signals remained below an exclusion threshold of 100 µV. Most of the movement related variance in the EEG activity is likely a movement artifact. Example of an EEG signal (black line) at movement onset that reached exclusion threshold (100 µV). Overlaid is the walking speed of the participant (green line). **(B)**As in **A**, but for an example of an EEG signal at movement onset that did not reach exclusion threshold (100 µV). (**C, D**) As in **A**, but for examples of EEG signals at movement onset with minimal movement contamination.

**Figure S2.**
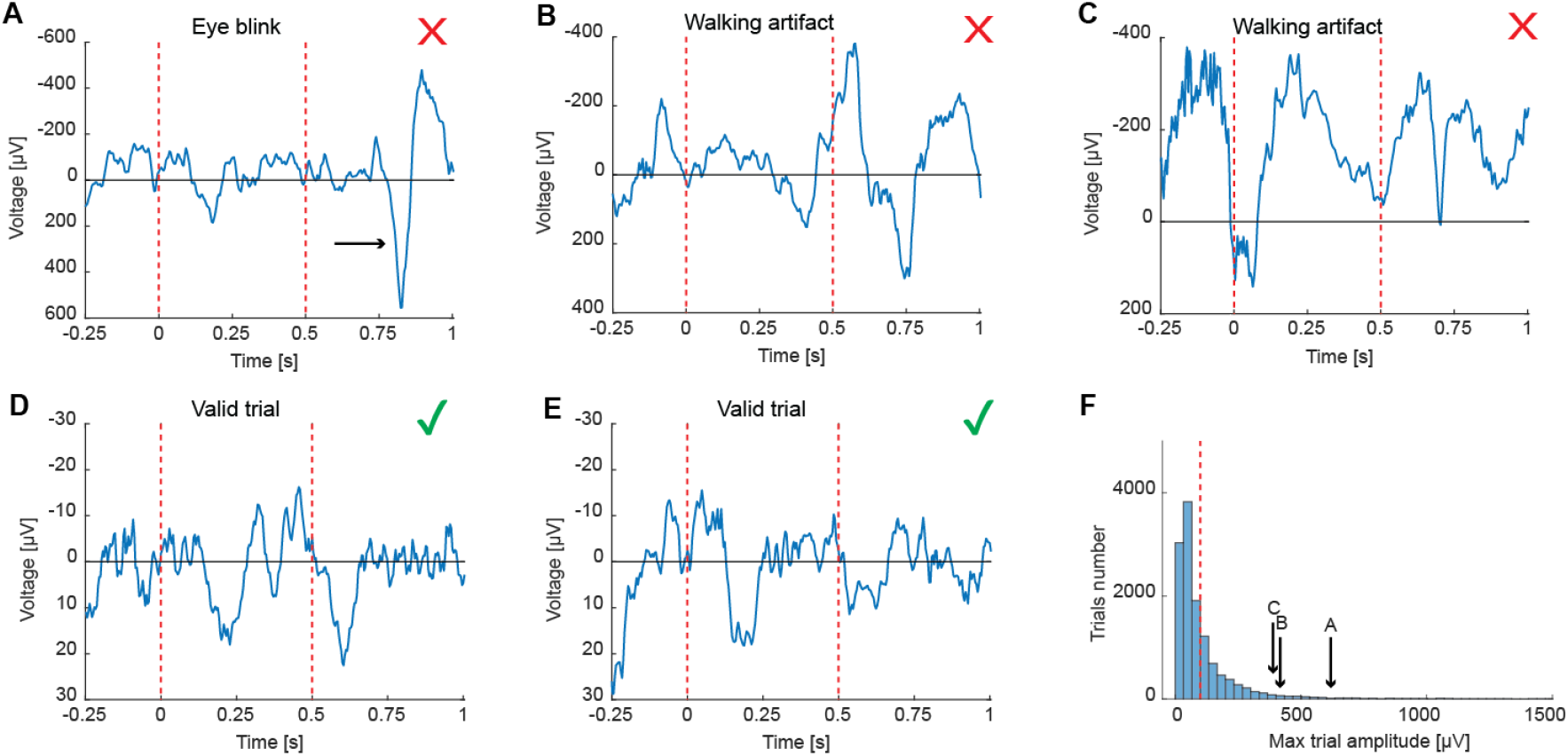
Examples of rejected and valid trials based on maximum signal amplitude in the visuomotor mismatch paradigm. (**A**) Example of an EEG response contaminated by an eye blink artifact (arrow). Dashed vertical red lines are onset and offset of the visuomotor mismatch. This trial was removed. (**B, C**) As in **A**, but for examples of EEG responses contaminated by walking artifacts. These trials were removed. (**D, E**) As in **A**, but for an example of an EEG response that met the inclusion criteria (amplitude < 100 µV). (**F**) Histogram of maximum trial amplitudes. The red dashed line marks the threshold for inclusion. 53 trials with amplitudes exceeding 1500 µV are not shown on the histogram.

**Figure S3.**
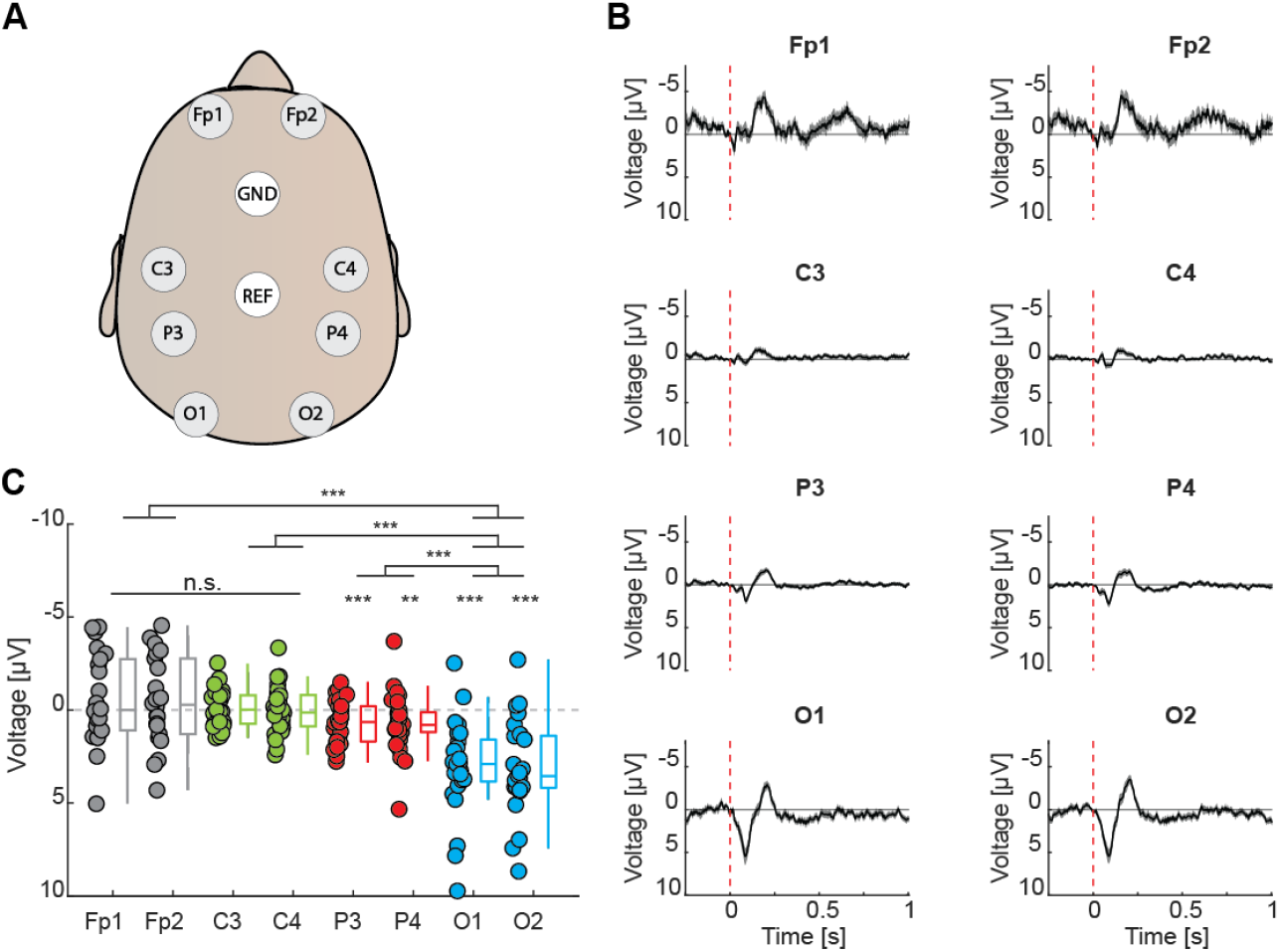
Visual responses are strongest over occipital cortex. **(A)**Top down view of EEG electrode locations on the head. **(B)**Visual evoked responses measured on electrodes shown in **A**. Solid black lines represent the mean, and shading indicates the SEM across participants. Dashed vertical red line is the onset of the checkerboard inversion. **(C)**Comparison of the response strength of visual evoked potentials measured on electrodes shown in **A**. Average response strength was calculated within a 100 ms window centered on the peak of the average visual response across all electrodes. Boxes mark median, quartiles, and range of data not considered outliers. Each circle represents data from an individual participant. ***: p<0.001, **: p<0.01, n.s.: not significant. See **Table S1** for all statistical information.

**Figure S4.**
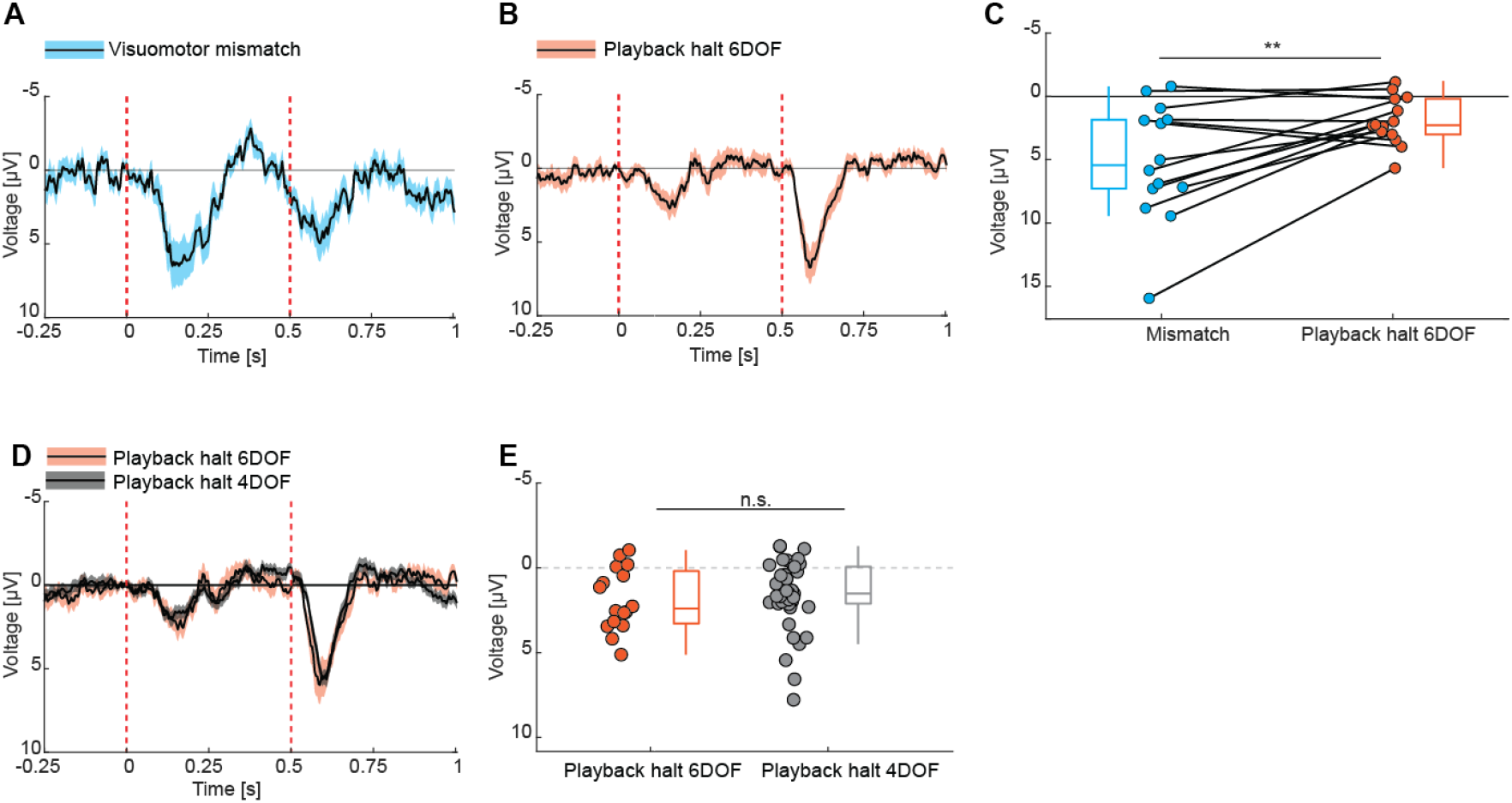
Visuomotor mismatch responses are bigger than playback halt responses even when the coupling is full. **(A)**Responses to visuomotor mismatches recorded from occipital electrodes. Solid black line represents the mean, and shading indicates the SEM across participants. Dashed vertical red lines are onset and offset of the visuomotor mismatch. **(B)**As in **A**, but showing playback halt responses to full six degrees of freedom (6DOF) visual flow playback, recorded over occipital electrodes **(C)**Comparison of the response strength to visuomotor mismatches and 6DOF playback halts. Average response strength was calculated within a 100 ms analysis window centered on the peak of the visuomotor mismatch response. Boxes mark median, quartiles, and range of data not considered outliers. Each data point corresponds to one participant and lines connect mismatch and playback halt responses from the same participant. **: p<0.01. See **Table S1** for all statistical information. **(D)**Comparison of 6DOF and 4DOF playback halt responses recorded from occipital electrodes. Solid black lines represent the mean, and shading indicates the SEM across participants. Dashed vertical red lines are onset and offset of the visuomotor mismatch. **(E)**Comparison of the response strength to 6DOF and 4DOF playback halts. Average response strength was calculated within a 100 ms analysis window centered on the peak of the playback halt response in the 6DOF condition. Each circle represents data from an individual recording session. Boxes mark median, quartiles, and range of data not considered outliers. n.s.: not significant.

**Figure S5.**
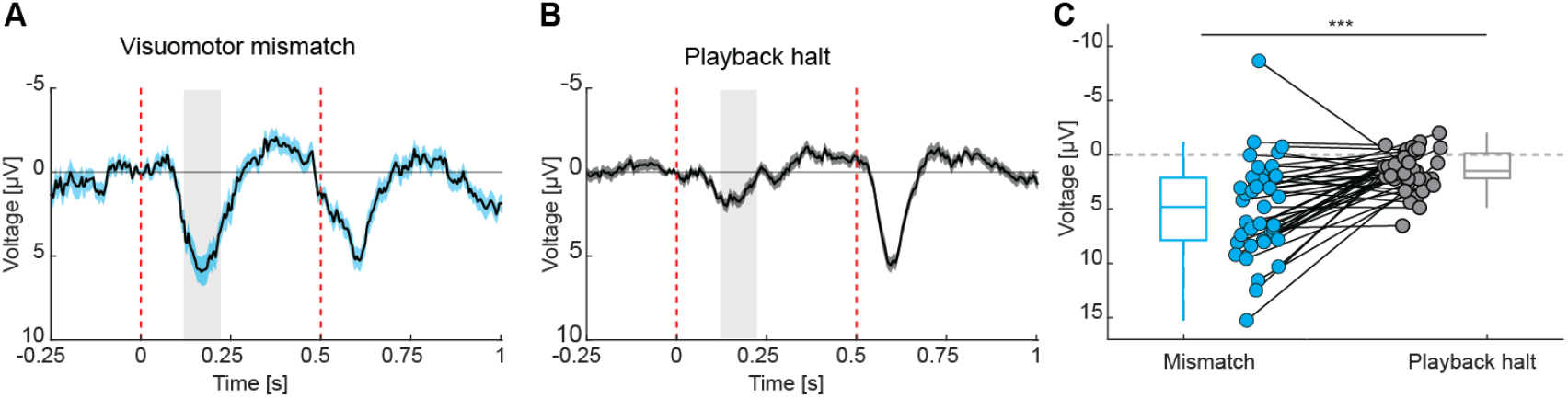
Mismatch and playback halt responses obtained from the same participants. **(A)**Responses to visuomotor mismatches recorded from occipital electrodes (O1 and O2). Solid black line represents the mean, and shading indicates the SEM across participants. The gray shaded areas mark the analysis windows used to quantify response strength in **C**. Dashed vertical red lines are onset and offset of the visuomotor mismatch. As in **Figure 2E, F**, but only including data from participants for which we have both closed and open loop data. **(B)**As in **A**, but for visual flow playback halt responses recorded from occipital electrodes. **(C)**Comparison of the response strength to visuomotor mismatch and playback halts (32 participants 4DOF and 7 participants 6DOF). Average response strength was calculated within a 100 ms analysis window centered on the peak of the visuomotor mismatch response. Boxes mark median, quartiles, and range of data not considered outliers. Each data point corresponds to one participant and lines connect mismatch and playback halt responses from the same participant. ***: p<0.001. See **Table S1** for all statistical information.

**Figure S6.**
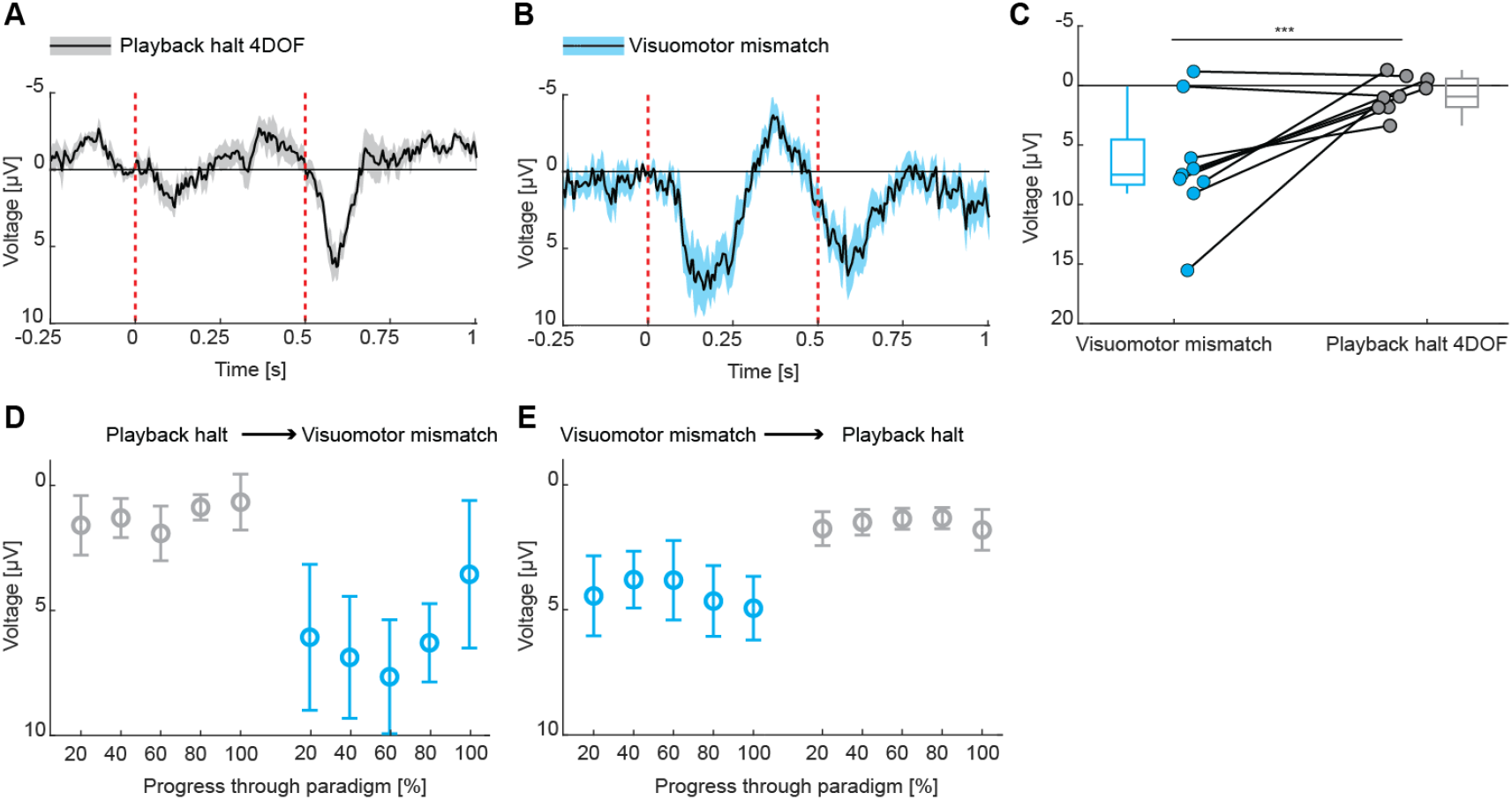
Adaptation over sessions does not explain the difference between visuomotor mismatch and playback halt responses. **(A)**Playback halt responses recorded from occipital electrodes in the reversed sequence experiment where the open loop session was presented first, followed by the closed loop session. The visual flow played back was the self-generated visual flow of the previous participants’ closed loop session. Note, this implies that visual flow statistics are no longer matched for closed and open loop for the same participant. Solid black line represents the mean, and shading indicates the SEM across participants. Dashed vertical red lines are onset and offset of the visual playback halt. **(B)**As in **A**, but for visuomotor mismatch responses. **(C)**Comparison of the response strength to visuomotor mismatches and playback halts in the reverse sequence session. Average response strength was calculated within a 100 ms analysis window centered on the peak of the visuomotor mismatch response. Boxes mark median, quartiles, and range of data not considered outliers. ***: p<0.001. See **Table S1** for all statistical information. **(D)**Playback halt and visuomotor mismatch responses as a function of progress through the session, when open loop session preceded closed loop session. Progress in the paradigm is measured as the percentage of total visuomotor mismatch or playback halt events. **(E)**As in **D**, but when closed loop was preceding the open loop session.

**Figure S7.**
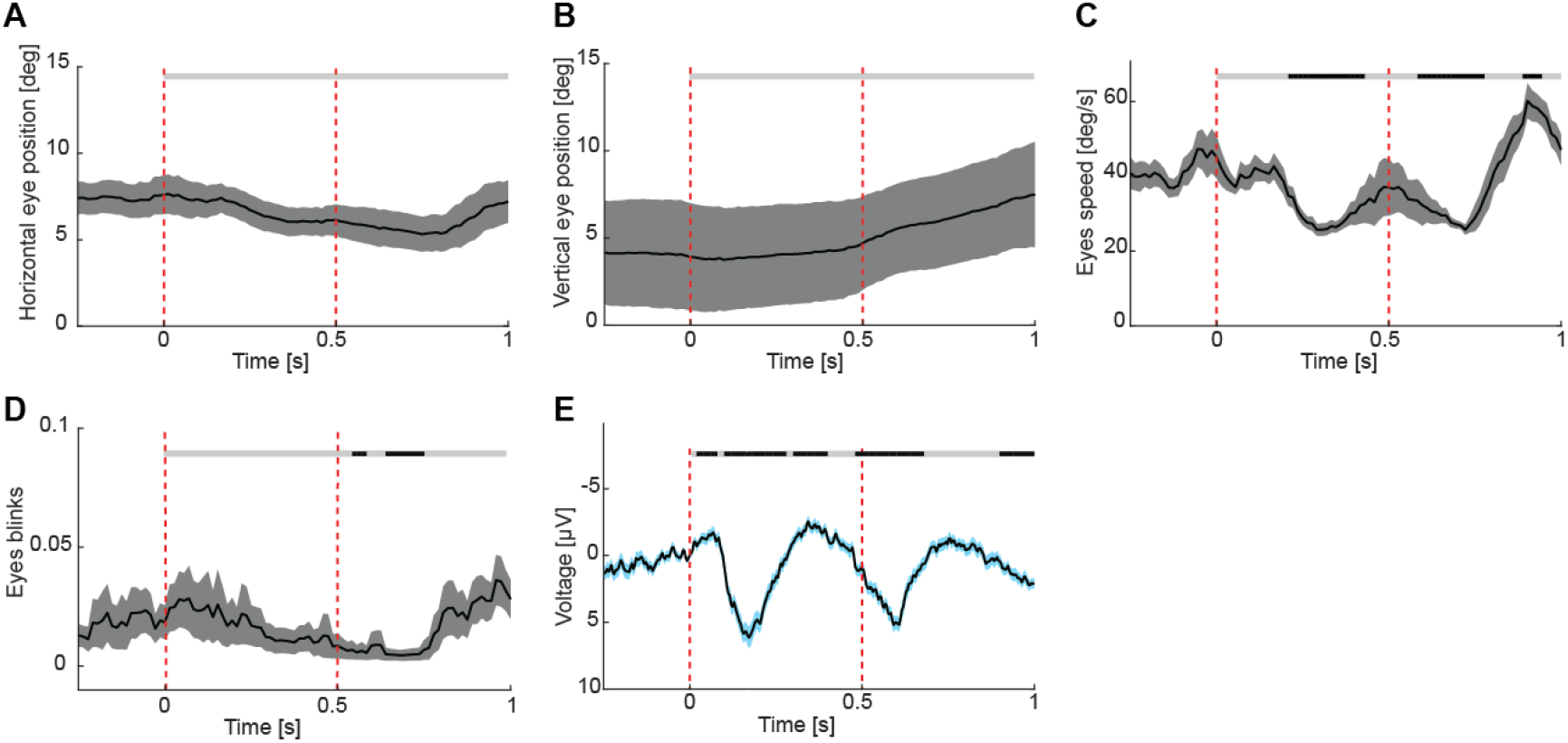
Visuomotor mismatch response cannot be explained by eye blinks or changes in eye movement speed. **(A)**Average horizontal eye position during visuomotor mismatch. Solid black line represents the mean, and shading indicates the SEM across participants. Vertical red dashed lines indicate mismatch onset and offset. The horizontal bar above the plot marks time bins in which the response differs significantly (black, p < 0.05) or not (gray, p > 0.05) from the baseline. **(B)**Average vertical eye position during visuomotor mismatch. **(C)**Average eye-movement speed during visuomotor mismatch. **(D)**Average probability of eye blinks during visuomotor mismatch. 0 – eyes open, 1 – eyes fully closed. **(E)**Responses to visuomotor mismatches recorded from occipital electrodes (O1 and O2).

**Figure S8.**
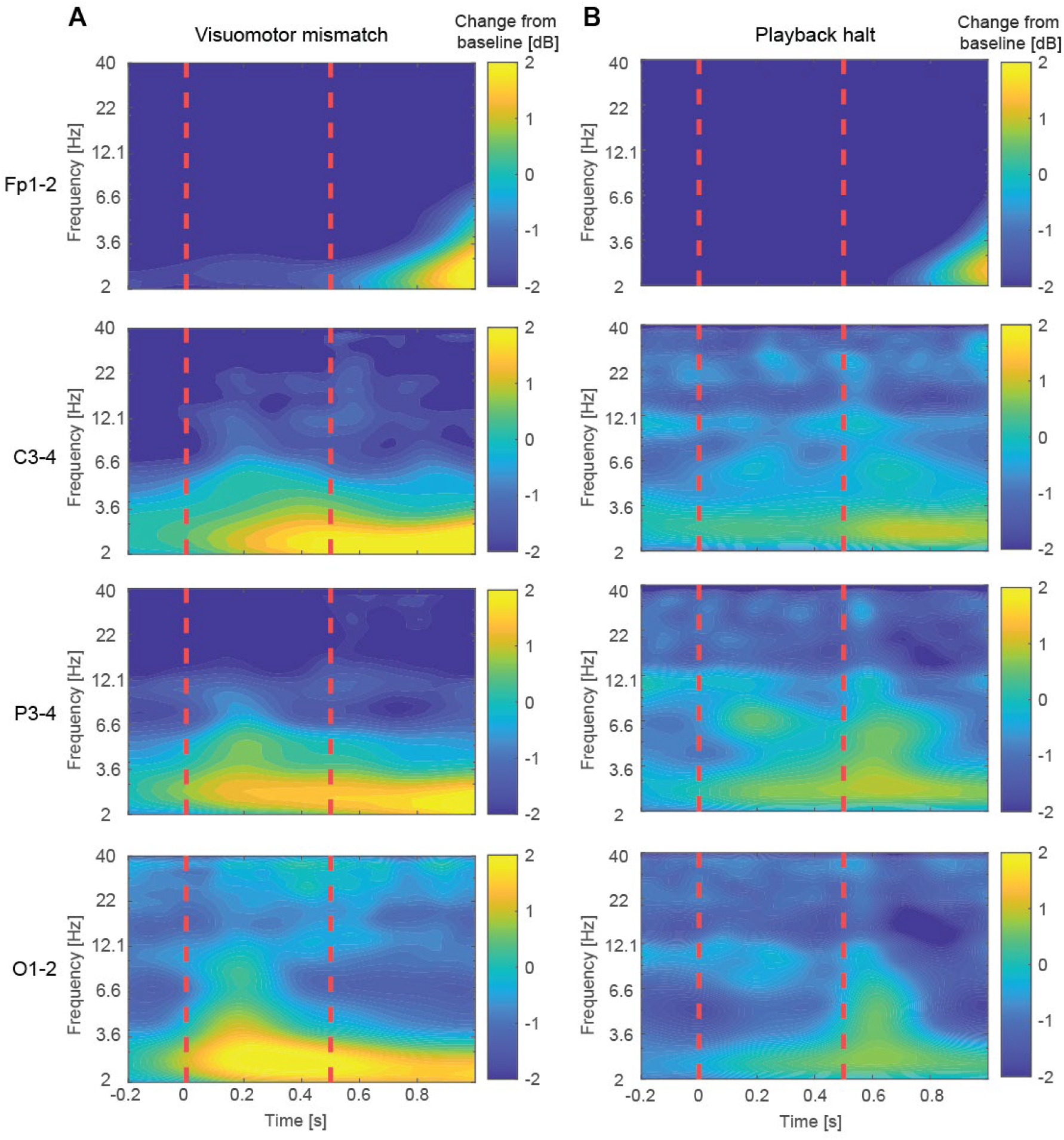
Time–frequency representations of EEG power changes during visuomotor mismatch and playback halt responses. **(A)**Time–frequency maps showing changes in spectral power relative to baseline for electrodes Fp1–2, C3–4, P3–4, and O1–2 following visuomotor mismatch presentation. Dashed vertical red lines are onset and offset of the visuomotor mismatch. **(B)**As in A, but for playback halts.

**Figure S9.**
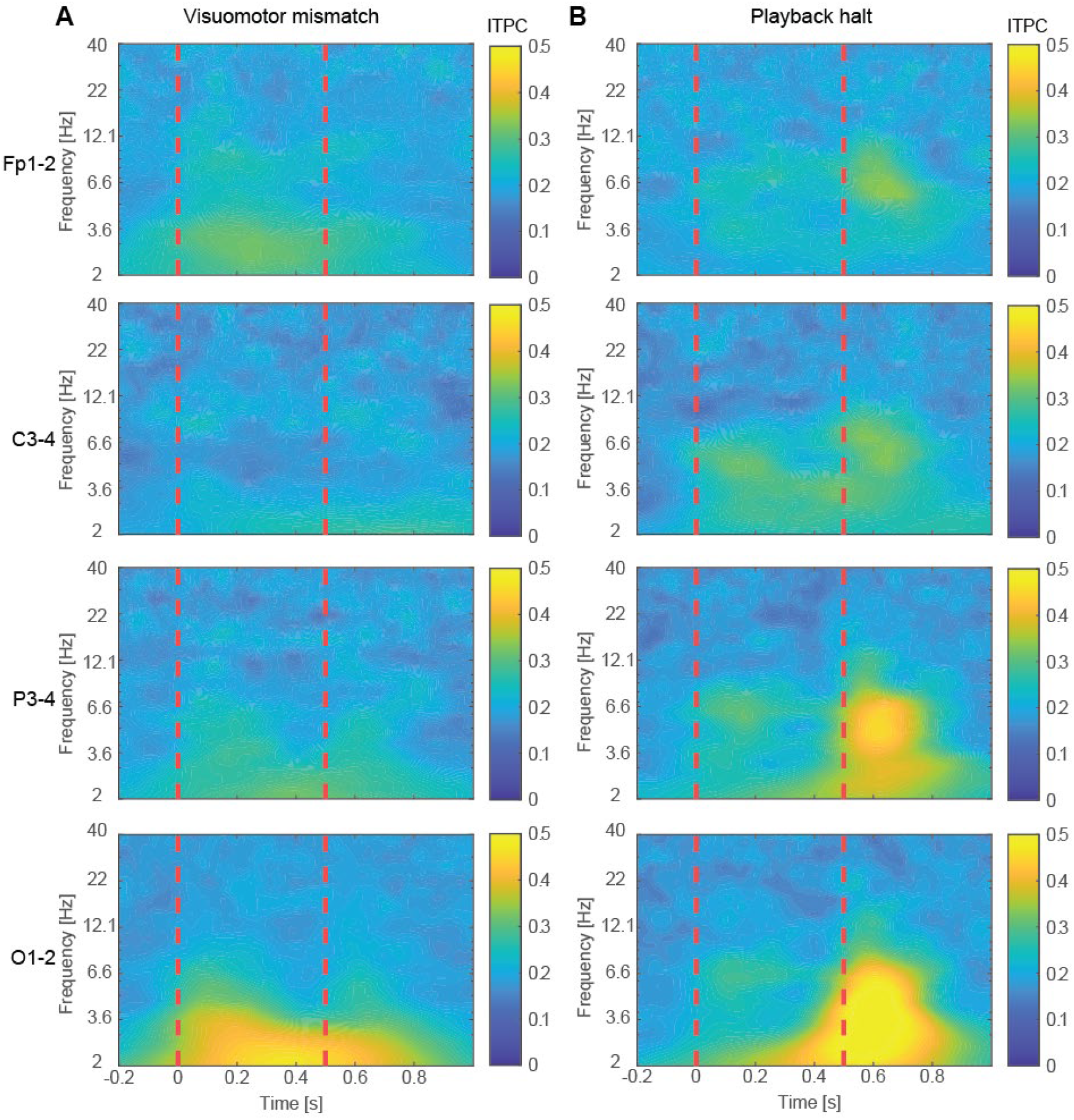
Inter-trial phase coherence (ITPC) for the visuomotor mismatch and playback halt responses. **(A)**ITPC across trials for electrode pairs Fp1–2, C3–4, P3–4, and O1–2 following visuomotor mismatch presentation. Dashed vertical red lines are onset and offset of the visuomotor mismatch. **(B)**As in A, but for playback halts.

**Figure S10.**
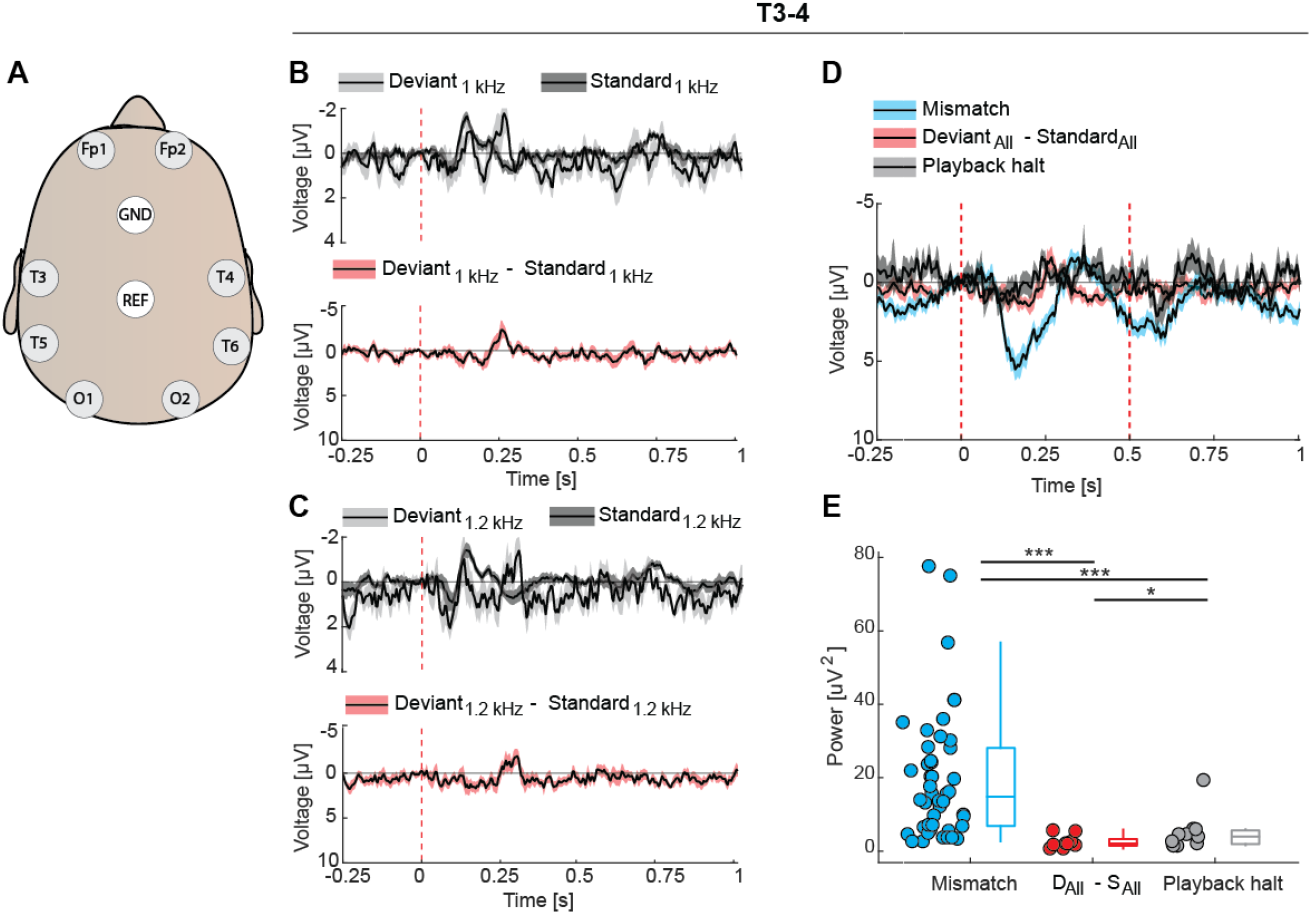
Visuomotor mismatch responses exceed auditory oddball mismatch responses at electrodes T3–T4. **(A)**Top-down view of EEG electrode positions on the scalp, including the temporal electrodes T3, T4, T5, T6. **(B)**Auditory responses to the 1 kHz tone presented as a standard versus as a deviant. Bottom: Oddball mismatch response calculated by subtracting the average response over trials in which the tone was presented as a standard from those when the tone was presented as a deviant. Solid black lines represent the mean, and shading indicates the SEM across participants. Dashed vertical red line is the onset of the auditory stimulus. **(C)**As in **B**, but for the 1.2 kHz tone. **(D)**Comparison of visuomotor mismatch and oddball mismatch responses (average over data shown in panels A and B) recorded from temporal electrodes. Solid black lines represent the mean response, with shading indicating SEM across electrodes. Dashed vertical red lines are onset (visual, mismatch) and offset (mismatch) of the stimuli. **(E)**Comparison of the power of visuomotor mismatch and oddball mismatch response (average over data shown in panels A and B), calculated within a 0 s - 0.5 s time window following stimulus onset. Boxes mark median, quartiles, and range of data not considered outliers. Each circle represents data from an individual participant. ***: p<0.001, *: p<0.05. See **Table S1** for all statistical information.

**Figure S11.**
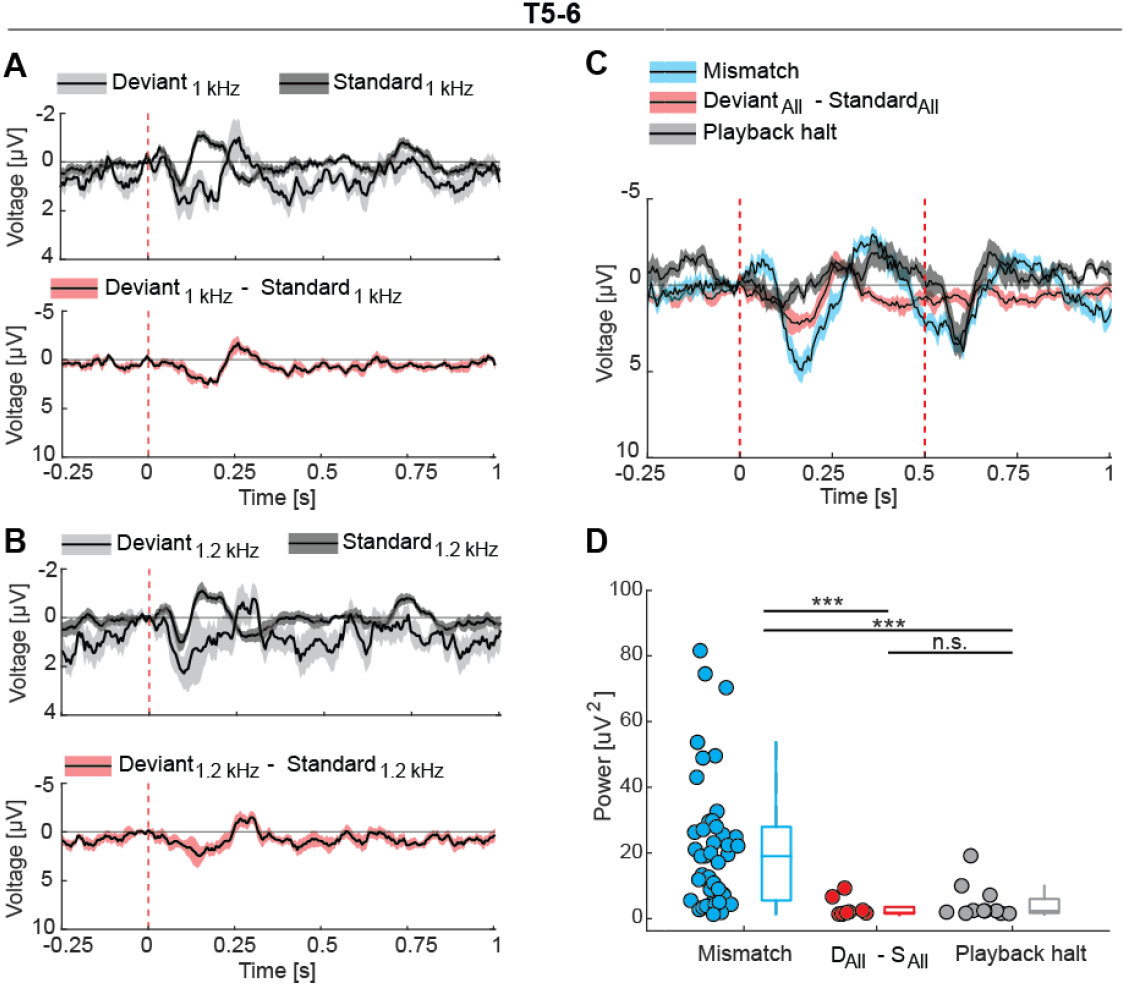
Visuomotor mismatch responses exceed auditory oddball mismatch responses at electrodes T5–T6. **(A)**Auditory responses to the 1 kHz tone presented as a standard versus as a deviant. Bottom: Oddball mismatch response calculated by subtracting the average response over trials in which the tone was presented as a standard from those when the tone was presented as a deviant. Solid black lines represent the mean, and shading indicates the SEM across participants. Dashed vertical red line is the onset of the auditory stimulus. **(B)**As in A, but for the 1.2 kHz tone. **(C)**Comparison of visuomotor mismatch and oddball mismatch responses (average over data shown in panels A and B) recorded from posterior temporal electrodes. Solid black lines represent the mean response, with shading indicating SEM across electrodes. Dashed vertical red lines are onset (visual, mismatch) and offset (mismatch) of the stimuli. **(D)**Comparison of the power of visuomotor mismatch and oddball mismatch response (average over data shown in panels A and B), calculated within a 0 s - 0.5 s time window following stimulus onset. Boxes mark median, quartiles, and range of data not considered outliers. Each circle represents data from an individual participant. ***: p<0.001, n.s.: not significant.

**Table S1.**
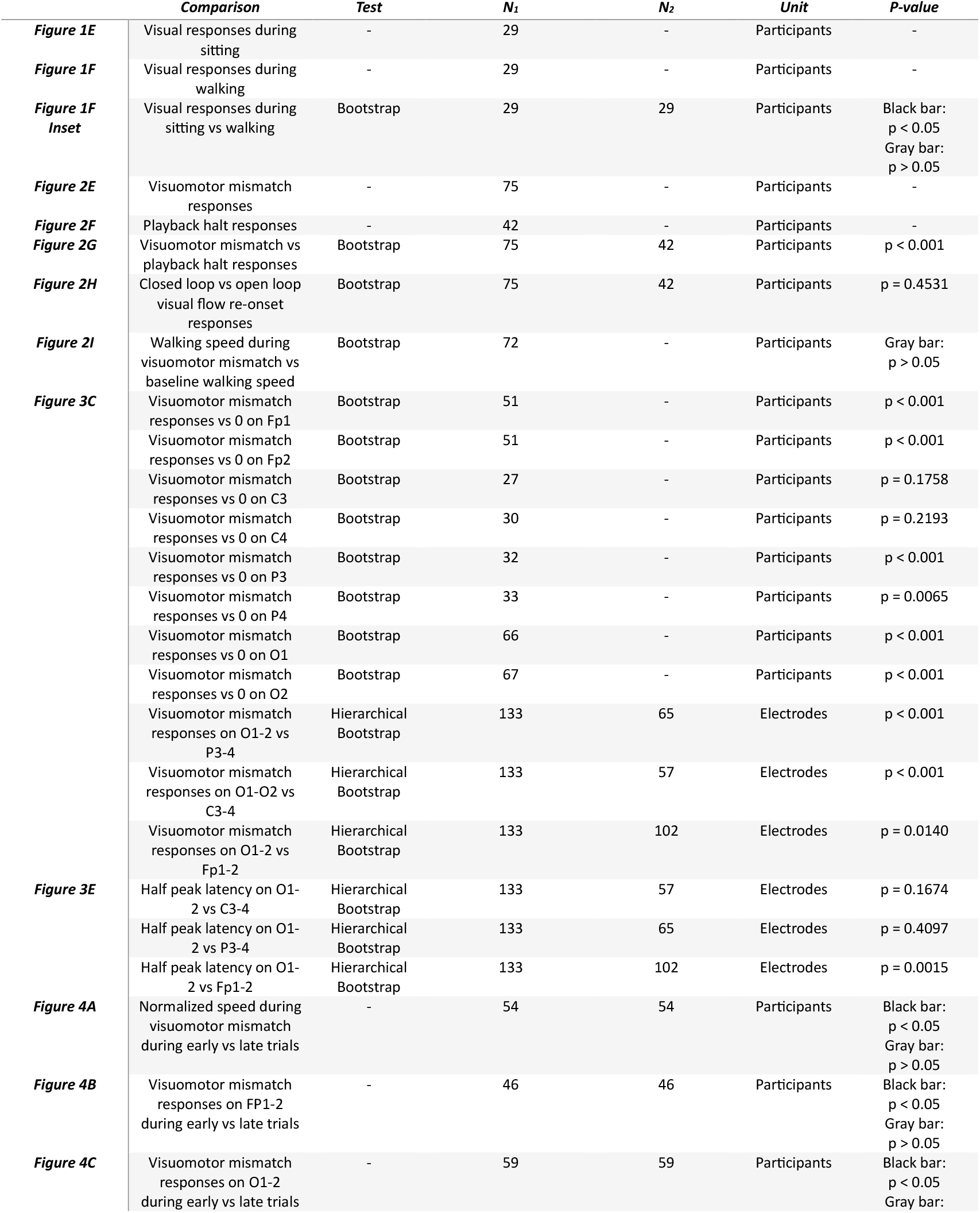
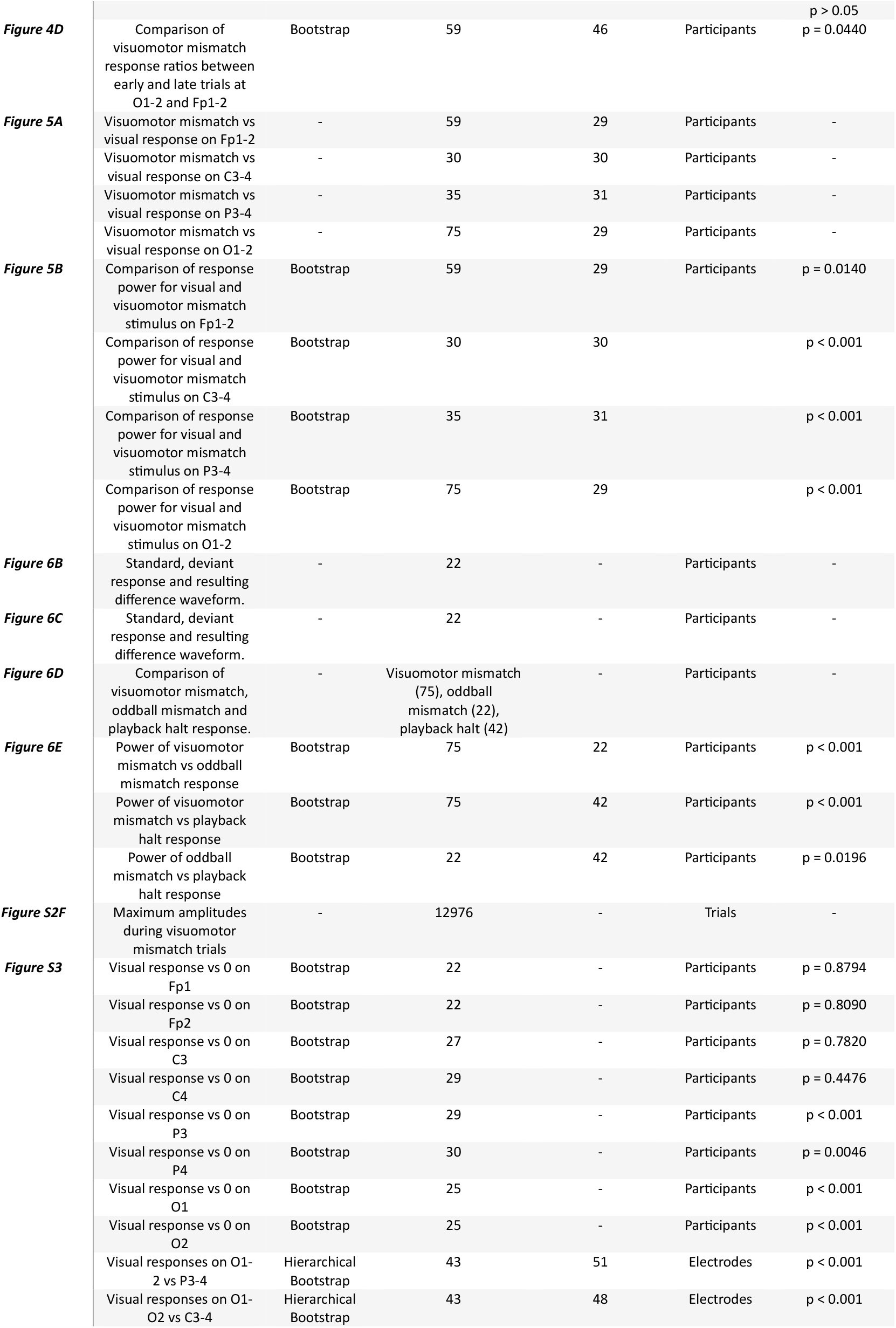
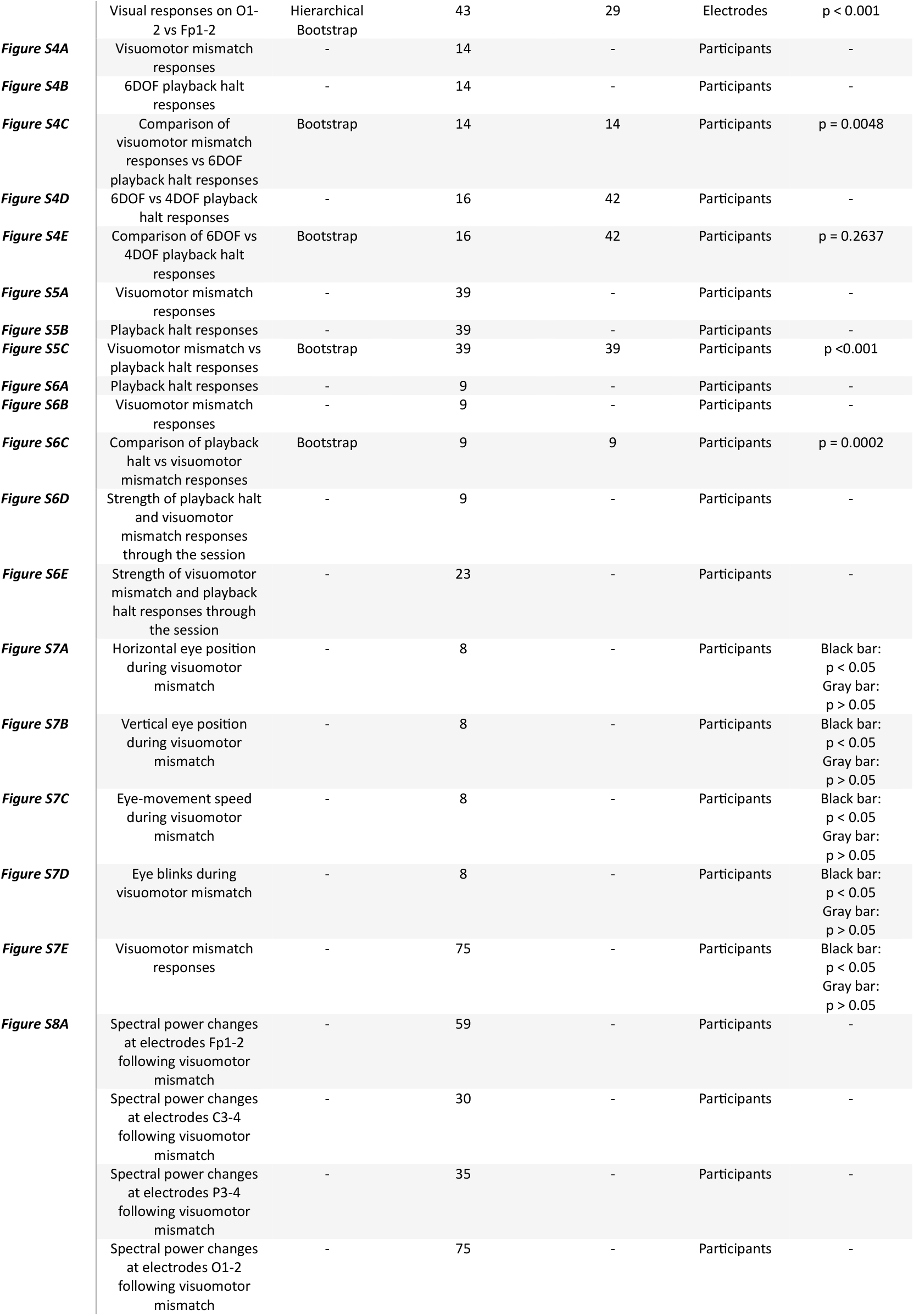
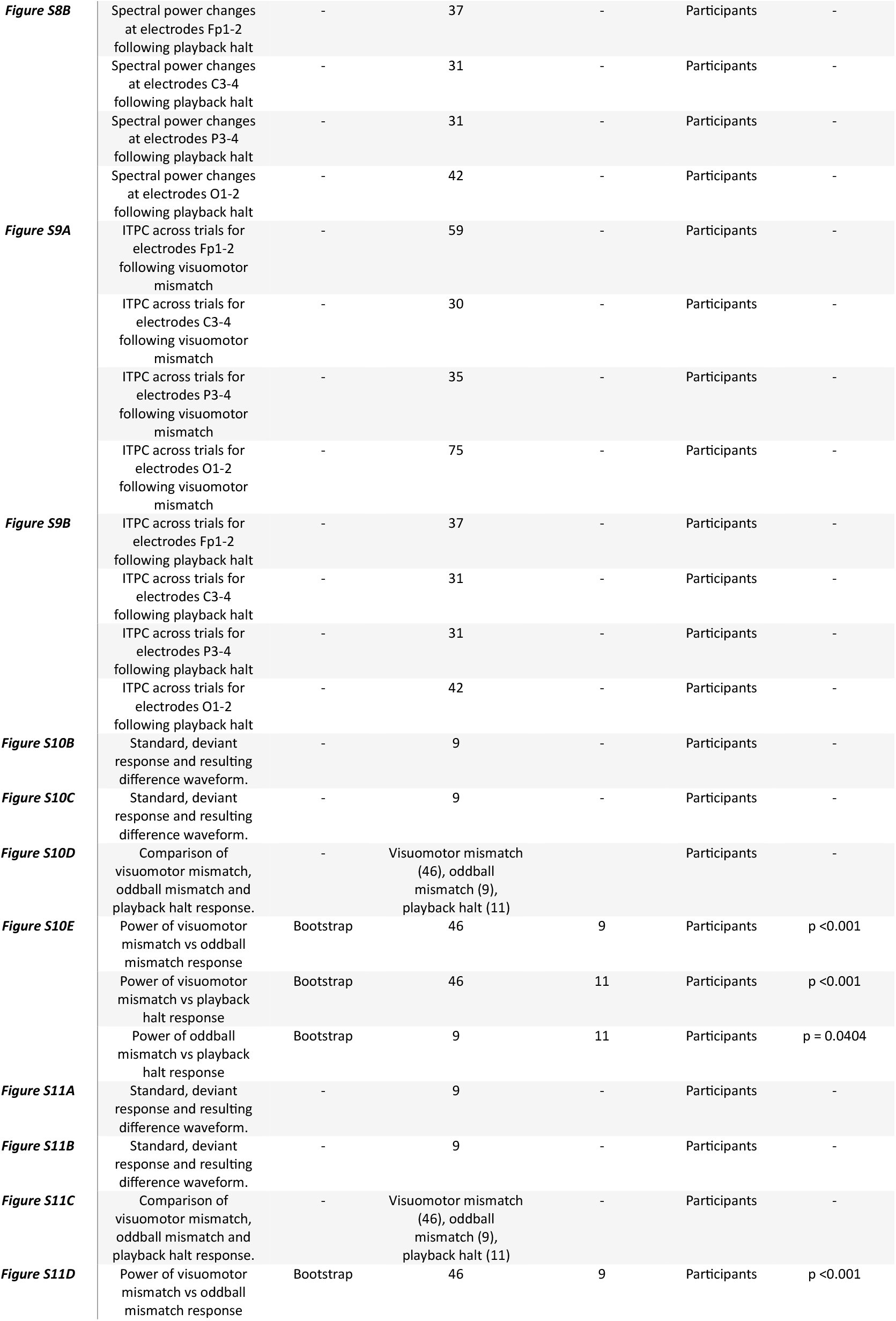
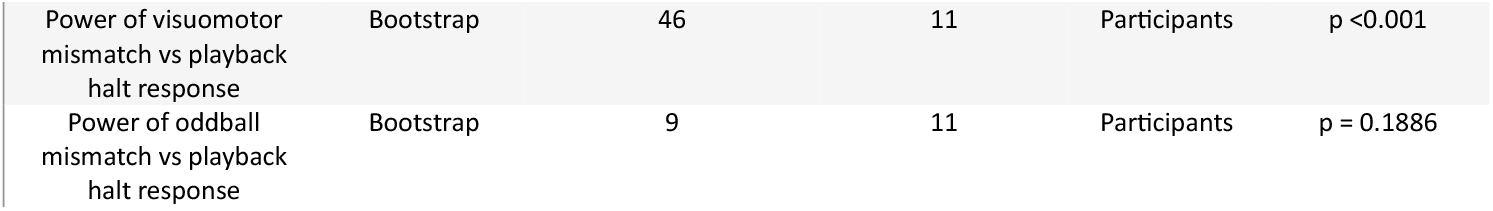
Statistics. We used hierarchical bootstrap (Saravanan et al., 2020) for all comparisons. We recorded a total of 91 sessions. The numbers in the table indicate the subset of these we could include for each analysis. Note, this differs for electrode location and condition. Exclusion reasons were a) recording is too noisy, or b) participant aborted the recording (in the case of open loop session).

**Table S2.** Excluded trials by paradigm and session. Indicated are the number and percentage of excluded trials as: excluded trials/all trials, % of excluded trials, for frontal (FP1-2), central (C3-4), parietal (P3-4), and occipital (O1-2) electrodes. See **Figure 3A** for electrode positions.

| PARADIGM | SESSION | Fp1-2 | C3-4 | P3-4 | O1-2 |
| --- | --- | --- | --- | --- | --- |
| VISUAL | Walking | 764/1610, **47.4%** | 428/1536, **27.9%** | 389/1536, **25.9%** | 597/1610, **37.1%** |
| VISUAL | Sitting | 379/1541, **24.6%** | 15/1455, **1.0%** | 14/1455, **1.0%** | 24/1541, **1.6%** |
| VISUOMOTOR | Closed loop (walking) | 971/2200, **44.1%** | 318/1044, **30.5%** | 211/1044, **20.2%** | 478/2200, **21.7%** |
| VISUOMOTOR | Open loop (sitting) | 241/1052, **22.9%** | 1/785, **0.1%** | 0/785, **0%** | 4/1052, **0.4%** |

## Key Resources Table

| Key Resources Table |  |  |  |  |
| --- | --- | --- | --- | --- |
| Reagent type (species) or resource | Designation | Source or reference | Identifiers | Additional information |
| Software, algorithm | MATLAB (2021b) | The MathWorks | RRID: SCR_001622 | Data analysis |
| Software, algorithm | Unity | Unity Technologies |  | Virtual reality design |
| Software, algorithm | Fusion 360 | Autodesk |  | Design of 3D objects for virtual reality corridors |
| Hardware, VR headset | Meta Quest 3, Meta Quest Pro | Meta |  | VR environment in EEG experiments |
| Hardware, EEG amplifier | OpenBCI Cyton Board | OpenBCI |  | 8 channel wireless EEG acquisition |
| Hardware, EEG electrode cap | EEG Electrode Cap Kit | OpenBCI |  | 19 channel electrode cap with sintered Ag/AgCl electrodes |
| Electrode Cap Gel | OpenBCI Electro-Gel | OpenBCI |  | Conductive gel used with EEG electrode caps for signal quality improvement. |

